# Targeting of scavenger receptors Stabilin-1 and Stabilin-2 ameliorates atherosclerosis by a plasma proteome switch mediating monocyte/macrophage suppression

**DOI:** 10.1101/2022.07.01.497917

**Authors:** Calin-Petru Manta, Thomas Leibing, Mirco Friedrich, Hendrik Nolte, Monica Adrian, Kai Schledzewski, Jessica Krzistetzko, Christof Kirkamm, Christian David Schmid, Yannick Xi, Ana Stojanovic, Sarah Tonack, Carolina de la Torre, Seddik Hammad, Stefan Offermanns, Marcus Krüger, Adelheid Cerwenka, Michael Platten, Sergij Goerdt, Cyrill Géraud

## Abstract

**Background:** Scavenger receptors (SR) Stabilin-1 (Stab1) and Stabilin-2 (Stab2) are preferentially expressed by liver sinusoidal endothelial cells. They mediate the clearance of circulating plasma molecules controlling distant organ homeostasis. Studies suggest that Stab1 and Stab2 may impact atherosclerosis. Although subsets of tissue macrophages also express Stab1, hematopoietic Stab1 deficiency does not modulate atherogenesis. Here, we comprehensively studied how targeting Stab1 and Stab2 affects atherosclerosis.

**Methods:** ApoE-KO mice were interbred with Stab1-KO and Stab2-KO mice and fed a Western diet (WD). For antibody targeting, Ldlr-KO mice were also used. Unbiased plasma proteomics were performed and independently confirmed. Ligand binding studies comprised GST-pull down and endocytosis assays. Plasma proteome effects on monocytes were studied by single cell RNA sequencing *in vivo*, and by gene expression analyses of Stabilin-ligand-stimulated and plasma-stimulated bone marrow-derived monocytes/macrophages (BMDM) *in-vitro*.

**Results:** Spontaneous and WD-associated atherogenesis was significantly reduced in ApoE-Stab1- and ApoE-Stab2-KO. Similarly, inhibition of Stab1 or Stab2 by monoclonal antibodies (mAB) significantly reduced WD-associated atherosclerosis in ApoE-KO and Ldlr-KO. While neither plasma lipid levels nor circulating immune cell numbers were decisively altered, plasma proteomics revealed a switch in the plasma proteome, consisting of 231 dysregulated proteins comparing Wildtype with Stab1/2 single and Stab1/2-double KO, and of 41 proteins comparing ApoE-, ApoE-Stab1- and ApoE-Stab2-KO. Among this broad spectrum of common, but also disparate SR ligand candidates, Periostin, Reelin and TGFBi, known to modulate atherosclerosis, were independently confirmed as novel circulating ligands of Stab1/2. scRNA-Seq of circulating myeloid cells of ApoE-, ApoE-Stab1- and ApoE-Stab2-KO showed transcriptomic alterations in patrolling (Ccr2^-^/Cx3cr1^++^/Ly6C^lo^) and inflammatory (Ccr2^+^/Cx3cr1^+^/Ly6C^hi^) monocytes including downregulation of pro-atherogenic transcription factor Egr1. In Wildtype BMDM, ligand exposure alone did not alter Egr1 expression *in-vitro*. However, exposure to plasma from ApoE-Stab1- and ApoE-Stab2-KO mice showed a reverted pro-atherogenic macrophage activation as compared to ApoE-KO plasma including downregulation of Egr1 *in-vitro*.

**Conclusions:** Inhibition of Stab1/Stab2 mediates an anti-inflammatory switch in the plasma proteome including direct Stabilin ligands. The altered plasma proteome suppresses both patrolling and inflammatory monocytes and, thus, systemically protects against atherogenesis. Altogether, anti-Stab1- and anti-Stab2-targeted therapies provide a novel approach for the future treatment of atherosclerosis.

**Clinical Perspective:** *What is new?:*

- Inhibition of evolutionary conserved class H scavenger receptors Stabilin-1 and Stabilin-2 reduces aortic plaque burden in preclinical models.
- Atheroprotection is mediated likely through downregulation on transcriptional factor Egr1 in monocytes by multifaceted plasma protein changes.
- Transforming growth factor, beta-induced (TGFBi), Periostin (POSTN) and Reelin (Reln) are novel ligands of Stabilin-1 and Stabilin-2 and are implicated in atherosclerosis development.

*What are the clinical implications?:*

- Monoclonal anti-Stab1- and anti-Stab2 antibodies provide a novel approach for the future treatment of atherosclerosis.
- In the future, the plasma proteome composition may serve as a predictive factor, biomarker or surrogate parameter for cardiovascular disease in patients.

## Introduction

Despite improvement by lipid-lowering therapies, atherosclerosis is still responsible for the majority of cardiovascular mortality indicating an urgent clinical need to identify novel target gene candidates. In this regard, scavenger receptors (SR) represent a class of candidate target proteins, as diverse family members such as SR B1, FcγRIII, FcγRIIb (CD32b) and CD36 have been shown to be involved in the development of atherosclerosis in hyperlipidemic ApoE-KO and Ldlr-KO mice ^1-6^.

Stabilin-1 (Stab1) and Stabilin-2 (Stab2) are conserved class H SR ^7, 8^ expressed by sinusoidal endothelial cells (ECs) in the liver (LSECs), lymph nodes, spleen and bone marrow ^8, 9^. Besides LSECs, Stab1 expression is found on certain subsets of macrophages, and it has been shown to be involved in immune responses during metastasis and tumor growth, liver fibrosis and sepsis ^10-13^. Stab1 ligands include malondialdehyde-LDL ^12^, SPARC, SI-CLP ^14^ and placental lactogen ^15^. Stab2 is the main receptor for hyaluronan (HA); blocking antibodies or genetic ablation of Stab2 inhibit HA uptake and lead to increased systemic HA levels ^9, 16, 17^. Besides HA, Stab2 also binds other glycosaminoglycans including heparin ^18^. It is also involved in antisense oligonucleotide pharmacokinetics and nanoparticle uptake, and it controls VWF-FVIII complex half-life and immunogenicity ^19-21^. Both Stabilins can bind acLDL, oxLDL, advanced glycation end products (AGE) and collagen propeptides ^22, 23^.

Due to their ligand profiles, including acLDL and oxLDL, which is known to alter macrophage responses in atherosclerosis ^24, 25^, it can be hypothesized that Stab1 and Stab2 may be involved in atherosclerosis ^26-29^. This is further supported by Stab1 expression in macrophages of human atheroma samples ^30^. However, expression of Stab1 in macrophages alone does not suffice to alter aortic atherosclerosis as we have recently shown that Ldlr-KO mice with Stab1 deficiency limited to the hematopoietic system do not show altered plaque development ^31^. On the other hand, Stab2 was identified as a candidate gene for Aath5, a genetic trait locus in mice for atherosclerosis of the aortic arch, reducing aortic plaque burden^32, 33^.

While Stab1-deficient (Stab1-KO) and Stab2-deficient (Stab2-KO) mice show a normal life span, Stab1 and Stab2 double-deficient mice (Stab-DKO) develop severe glomerulosclerosis associated with albuminuria and mild to moderate sinusoidal liver fibrosis, and they exhibit a strongly reduced lifespan. By kidney transplantation, it was shown by us that indeed circulating blood factors are responsible for glomerulosclerosis in Stab-DKO mice, providing proof of principle that Stabilin-mediated clearance remotely controls distant organ homeostasis ^16^.

To assess the role of Stab1 and Stab2 in atherosclerosis *in-vivo*, ApoE-deficient (ApoE-KO) mice were interbred with Stab1-KO and Stab2-KO mice to investigate aortic plaque development. As models of atherosclerotic disease progression, we studied spontaneous plaque development over a period of 12 months and Western diet (WD) feeding for two months. To test whether treatment with mAB against Stab1 or Stab2, respectively, could recapitulate the effects in the genetic models, we treated Western diet-fed ApoE-KO and Ldlr-KO mice with anti-Stab1 and anti-Stab2 mAB. We show here that genetic deficiency of Stab1 or Stab2 as well as antibody targeting of either Stab1 or Stab2 significantly reduced aortic plaque development in spontaneous as well as WD-induced models of atherosclerosis. Mechanistically, we present evidence that reduction of atherosclerosis is mediated by protective factors in blood plasma of Stab1- and Stab2-deficient mice that revert pro-atherogenic monocyte/macrophage activation including suppression of patrolling and inflammatory monocytes.

## Methods

The data, methods, and study materials used to conduct the research will be available from the corresponding author on reasonable request.

The animal welfare commission of the Regierungspräsidium Karlsruhe, Germany approved all animal experiments. A detailed description of the methods and supporting data are available in the Data Supplement.

### Statistical Analysis

Statistical analyses, excluding scRNASeq and unbiased proteomics, were performed with JMP^®^ 14 and SAS^®^ 9.4M7 (SAS Institute Inc.). Data are presented as mean±SEM unless otherwise indicated. The used statistical tests are indicated in the Data Supplement. For statistical procedures regarding scRNASeq and unbiased proteomics, a detailed description is available in the Data Supplement.

## Results

### Genetic deficiency and antibody-mediated inhibition of Stab1 and Stab2 protect against progression of atherosclerosis in ApoE-KO and Ldlr-KO mice

To assess the role of Stab1 and Stab2 during atherosclerosis, murine models of aortic atherosclerosis were analyzed in mice with combined genetic deficiencies of ApoE and Stab1 (ApoE-Stab1-KO) and ApoE and Stab2 (ApoE-Stab2-KO). In comparison to ApoE-KO, ApoE-Stab1-KO and ApoE-Stab2-KO mice displayed significantly decreased spontaneous aortic plaque development at 12 months of age, as assessed by total plaque area in relation to total aortic area employing Oil-red O (ORO) staining and bright field microscopy (Fig. 1A). These findings were confirmed by assessing plaque abundance in histological cross sections of the aorta (Fig. 1B). Furthermore, ApoE-Stab1-KO and ApoE-Stab2-KO male mice also exhibited significantly decreased aortic plaque development after 2 months of WD-feeding in comparison to ApoE-KO mice (Fig. 1C).

**Figure 1.**
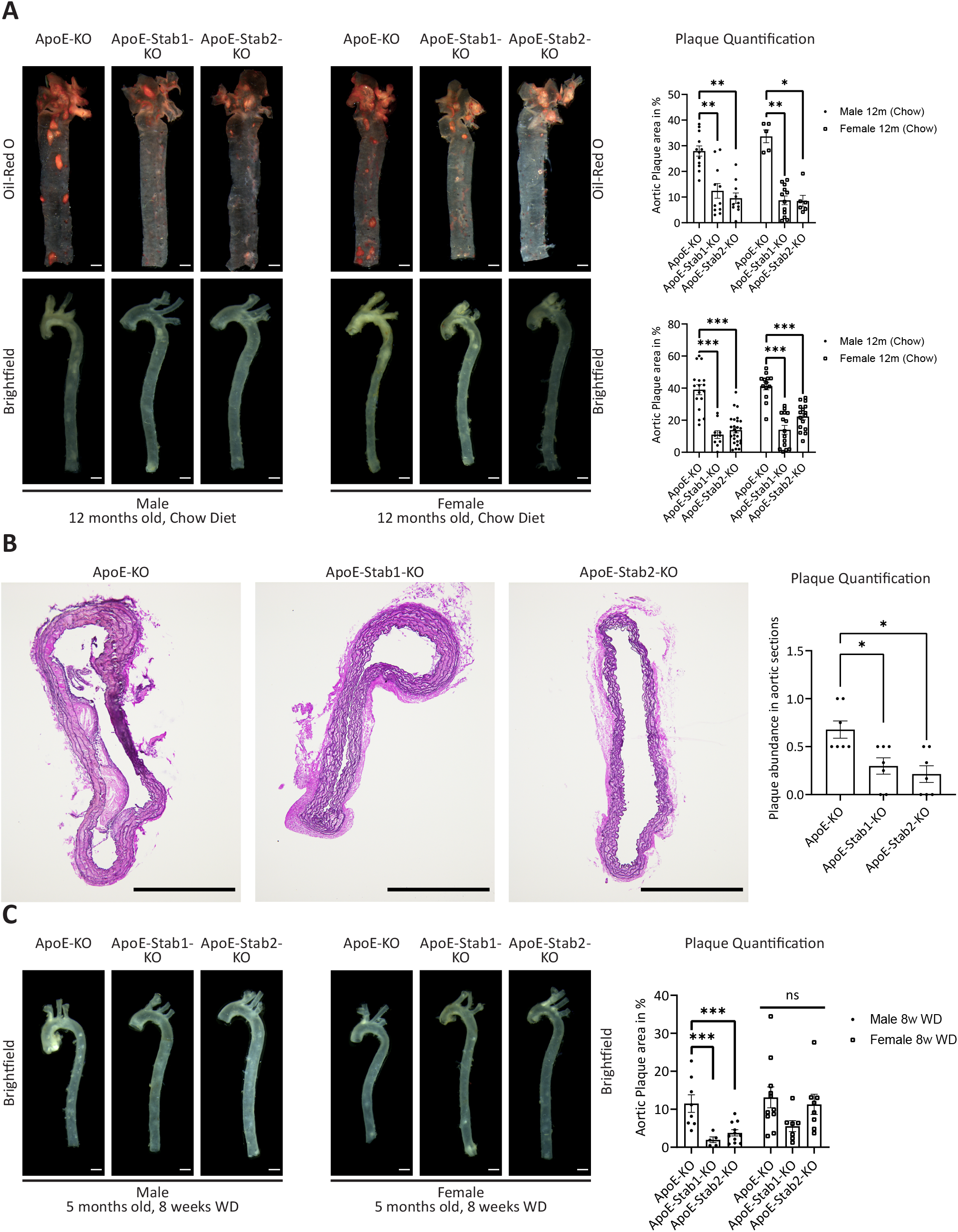
Decreased aortic plaque formation in ApoE-KO with genetic Stabilin-deficiency **(A and B)** Aortae of 12 months old ApoE-KO, ApoE-Stab1-KO and ApoE-Stab2-KO mice on chow diet. **(A)** Representative microscopic photographs of ORO staining (upper panel, n≥5) and unstained brightfield microscopy (lower panel, n≥10) of male (left) and female (right) mouse aortae. Quantification is shown on the right. Scale bar (white) = 500μm. **(B)** Representative Elastica-van-Gieson stained cross sections of aortae. Plaque abundance was evaluated on at least three (arch, thoracal and abdominal) sections per Aorta. Sections containing any plaque were counted and divided by the number of total sections per replicate. Scale bar (black) = 500μm, n=7. **(C)** Aortae of 5 months old ApoE-KO, ApoE-Stab1-KO and ApoE-Stab2-KO mice after 8 weeks of Western Diet (WD) feeding. Representative microscopic photographs of unstained brightfield microscopy of male (left panel) and female (right panel) aortae. Quantification is shown on the right. Scale bar (white) = 500μm, n≥5. Error bars represent SEM. ns=not significant, *p<0.05, **p=0.01, ***p<0.001.

As genetic deficiency reduced spontaneous and WD-associated plaque development in ApoE-KO mice, we were interested to know whether therapeutic inhibition of Stab1 and Stab2 might have similar effects. Therefore, ApoE-KO and Ldlr-KO mice were treated with mAB against Stab1 (anti-Stab1) and Stab2 (anti-Stab2) or with the corresponding isotype controls three times per week, while they were concomitantly fed a WD for 8 weeks or 12 weeks, respectively. Plaque development was assessed by analysis of brightfield and ORO-stained whole mounts as well by ORO-stained cross sections of aortic roots. Anti-Stab1 treatment significantly decreased plaque development in female ApoE-KO mice, while anti-Stab2 treatment showed a trend towards a significant reduction of plaque development in female ApoE-KO mice (Fig. 2A). In WD-fed, female Ldlr-KO mice, treatment with either anti-Stab1 or anti-Stab2 mAB for 12 weeks significantly reduced plaque burden in comparison to isotype control treatment (Fig. 2B). Although total plaque area was reduced, plaque composition in ApoE-KO and Ldlr-KO mice was not altered in regard to smooth muscle cell, total immune cell and macrophage content when comparing anti-Stab1, anti-Stab2 and isotype control treatment (Fig. 3A). Thus, genetic deficiency as well as antibody-mediated inhibition of Stab1 or Stab2 significantly decreased plaque development in different models of atherosclerosis development.

**Figure 2.**
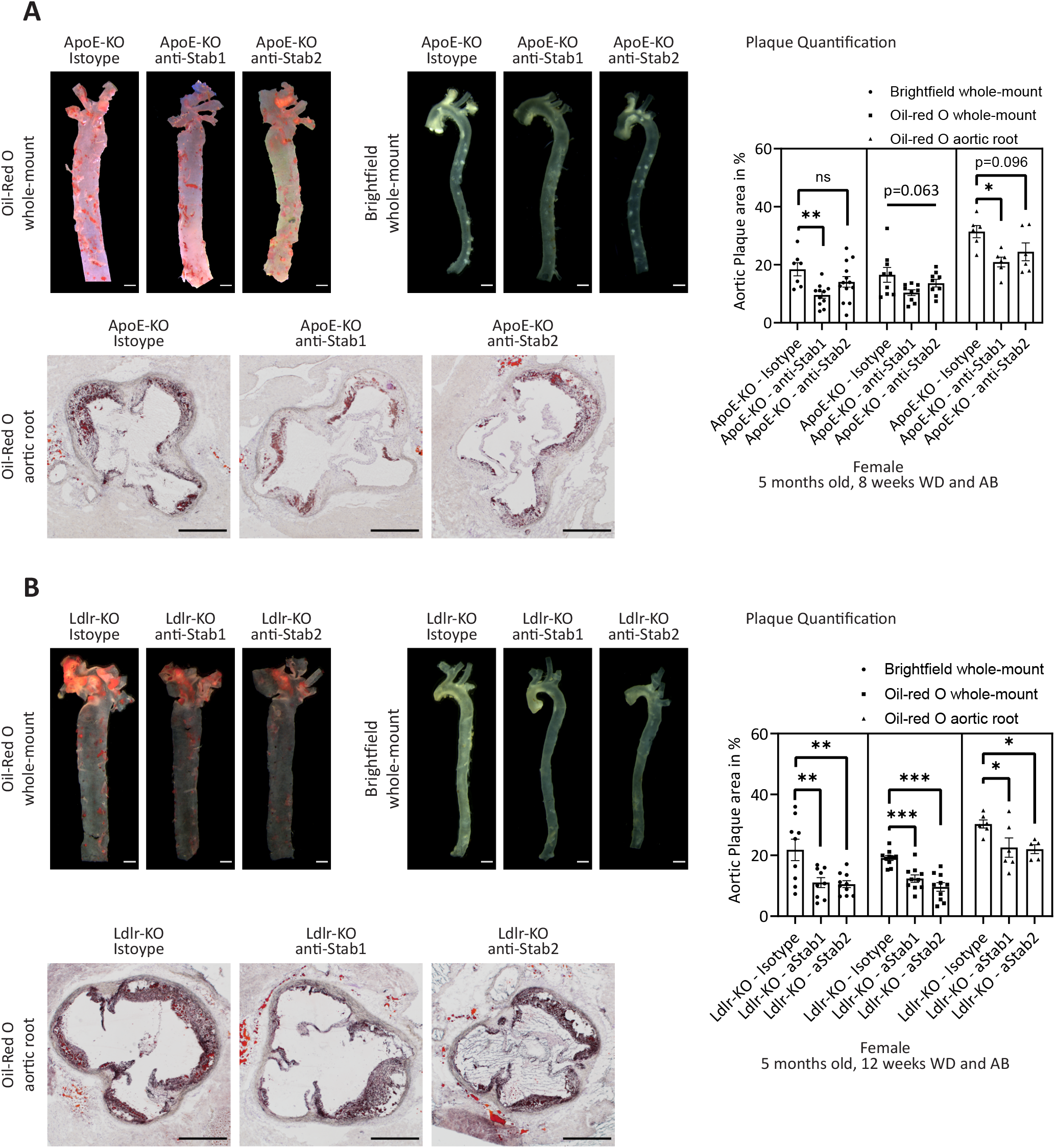
anti-Stabilin-antibodies reduce aortic plaque development in ApoE-KO and Ldlr-KO **(A)** Aortae of 5 months old ApoE-KO female mice after 8 weeks of Western Diet (WD) feeding and concomitant treatment with Isotype control antibody (Isotype), anti-Stab1 antibody (anti-Stab1) and anti-Stab2 antibody (anti-Stab2). Representative microscopic photographs of ORO staining (upper left panel, n=6), unstained brightfield microscopy (upper right panel, n≥7) of whole mouse aortae. Lower panel: ORO staining of aortic root cross sections (n=6). Quantification is shown on the right. Scale bar = 500μm. **(B)** Aortae of 5 months old female Ldlr-KO mice after 12 weeks of Western Diet (WD) feeding and concomitant treatment with Isotype, anti-Stab1 and anti-Stab2. Representative microscopic photographs of ORO staining (left panel, n≥9) and unstained brightfield microscopy (right panel, n=10). Quantification is shown on the right. Lower panel: ORO staining of aortic root cross sections (n≥5). Quantification is shown on the right. Scale bar (white) = 500μm. Error bars represent SEM. ns=not significant, *p<0.05, **p<0.01, ***p<0.001.

**Figure 3.**
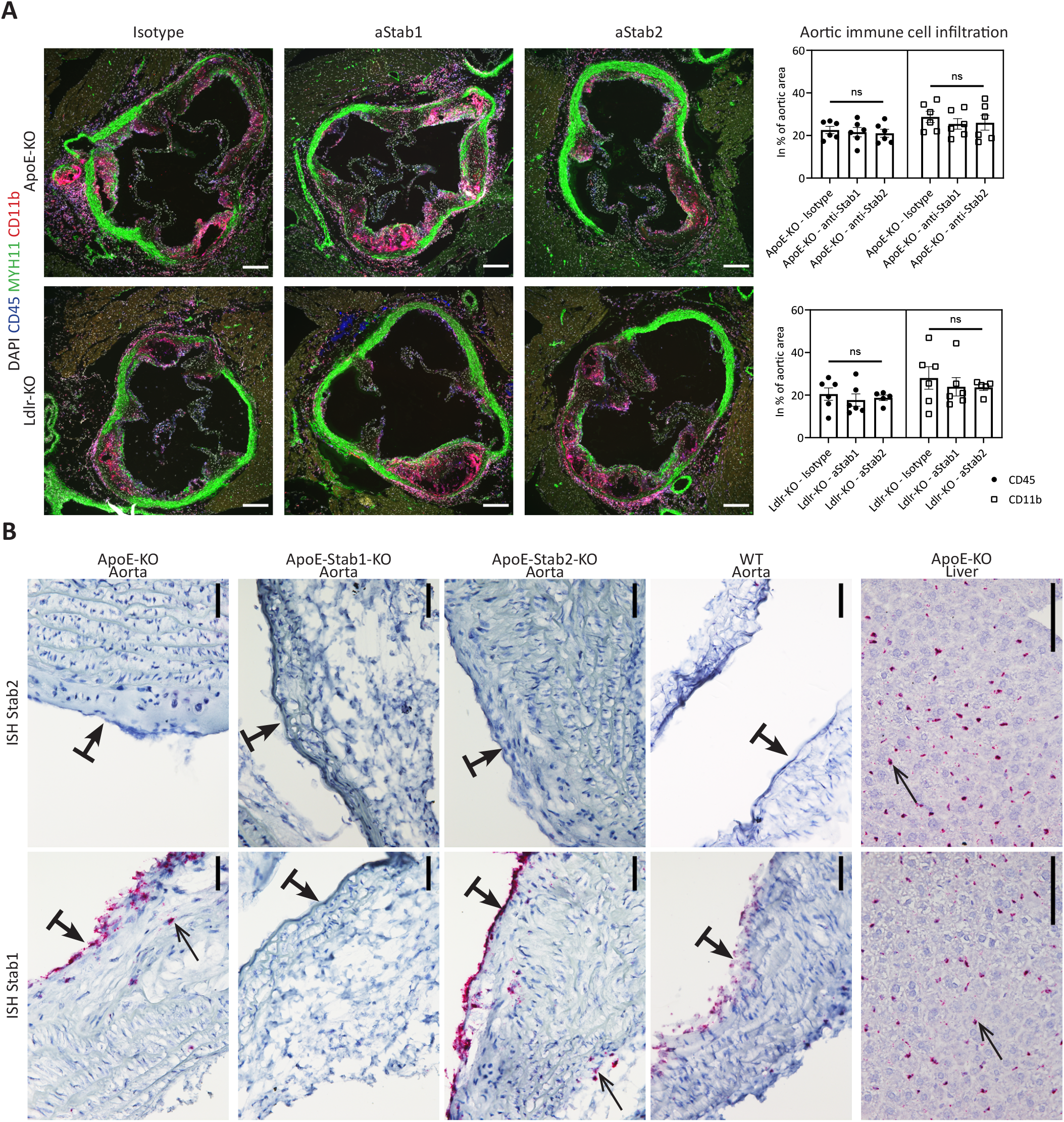
Aortic immune cell infiltration and expression of Stabilins in aortic plaques **(A)** Immunofluorescent co-staining with DAPI (white), Myosin-11 (MYH11, green), CD45 (blue) and CD11b (red) was performed on aortic root sections of ApoE-KO mice (upper panel, n≥6) treated with Stabilin mABs or Isotype control antibody or Ldlr-KO (lower panel, n≥5) mice treated with Stabilin mABs or Isotype control antibody. Scale bar (white) = 200μm. Quantification of CD45 and CD11b as percentage of aortic area is shown on the right. **(B)** In-situ hybridizations for Stab1 (lower panel) and Stab2 (upper panel) of aortae (left panels) of ApoE-KO, ApoE-Stab1-KO, ApoE-Stab2-KO and WT as well as hepatic tissues (right panel) of ApoE-KO as positive control (n≥3). Bold underlined arrows (**↥**) indicate endothelial lining. Bar is directed towards luminal side. Arrows (↑) indicate positive staining. Scale bar (black) = 100μm. Error bars represent SEM. ns=not significant, *p<0.05, **p<0.01, ***p<0.001.

Overall, treatment of mice with anti-Stab1 or anti-Stab2 mAB did not show major side effects while providing inhibition of Stabilin-mediated scavenging functions and protection from aortic plaque development. Female ApoE- and Ldlr-KO mice, treated with anti-Stab1 or anti-Stab2 mAB or isotype control antibodies for 8 (ApoE-KO) or 12 weeks (Ldlr-KO) showed comparable weight curves (Suppl. Fig. 2A). Histologic assessment of kidneys and livers of antibody-treated Ldlr-KO mice did not show signs of renal glomerulosclerosis or liver fibrosis (Suppl. Fig. 2B). Plasma levels of HA were significantly increased in both ApoE-KO and Ldlr-KO mice treated with anti-Stab2 mAB, but not with anti-Stab1 mAB, confirming the ability of anti-Stab1 or anti-Stab2 mAB to specifically interfere with the selective scavenging function of either of the two SR (Suppl. Fig. 2C).

### Stab1 and Stab2 are positioned to exert systemic control of aortic atherosclerosis, but do not strongly affect plasma lipid levels

Stab1 and Stab2 are expressed by LSECs, as well as by sinusoidal endothelial cells in lymph nodes, bone marrow and spleen. Stab1 is also known to be expressed by some tissue resident macrophages as well as subpopulations of M2-like monocytes/macrophages. Therefore, Stab1 and Stab2 exert local and systemic effects on aortic atherosclerosis. To assess potential local effects, we assessed expression of Stab1 and Stab2 in aortic tissue of WT, ApoE-KO, ApoE-Stab1-KO and ApoE-Stab2-KO mice by in situ hybridization (RNA-ISH). As expected, RNA-ISH did not reveal expression of Stab2 mRNA in aortic EC or other cells of aortic plaques, while showing strong expression in the liver (Fig. 3B). On the contrary, Stab1 mRNA was detected in aortic EC and some stromal plaque cells, likely macrophages, in ApoE-KO and ApoE-Stab2-KO mice. No signal for Stab1 was detected in ApoE-Stab1-KO mice (Fig. 3B). Therefore, Stab2 controls atherogenesis *via* its systemic clearance function while Stab1 may affect atherosclerosis *via* systemic as well as local mechanisms. As already mentioned, however, deficiency of Stab1 in hematopoietic stem cell-derived cells including macrophages alone is not sufficient to protect from atherosclerosis ^34^.

As Stab1 and Stab2 have been reported to bind certain lipids such as acLDL and oxLDL, plasma lipid profiles were assessed. In contrast to aortic plaque development, plasma lipid profiles were not consistently altered by genetic deficiency of Stab1 and Stab2 or by treatment with anti-Stab1 or anti-Stab2 mAB. Total cholesterol was slightly increased in WD-fed ApoE-Stab1-KO mice and in ApoE-KO mice treated with anti-Stab1, while LDL cholesterol was only significantly increased in ApoE-KO mice treated with anti-Stab1 (Suppl. Fig. 3). In the remaining groups, total cholesterol and LDL cholesterol were not significantly altered and HDL cholesterol, oxLDL and triglycerides did not show any significant differences among all models (Suppl. Fig. 3A). To assess whether Stabilin-dependent plasma changes control foam cell formation, RAW264.7 cells were exposed to plasma from ApoE-KO, ApoE-Stab1-KO and ApoE-Stab2-KO mice *in-vitro* and oxLDL uptake was quantified. Here, significant differences in oxLDL uptakes could not be detected (Suppl. Fig. 3B). Therefore, plasma lipid alterations cannot explain the alterations of aortic plaque development induced by genetic deficiency of Stab1 and Stab2 or by treatment with anti-Stab1 or anti-Stab2 mAB.

### Stab1 and Stab2 determine plasma protein composition

Stab1 and Stab2 act as endothelial SR that control the levels of a wide range of ligands in the peripheral circulation. However, the ligand repertoire of Stab1 and Stab2 has not yet been comprehensively characterized. As we hypothesized that Stab1 and Stab2 exert their effects on atherosclerosis via circulating ligands, we analyzed the plasma proteome of ApoE-KO, ApoE-Stab1-KO and ApoE-Stab2-KO mice (Fig. 4, Suppl. Data 1) as well as of WT, Stab1-KO, Stab2-KO and Stab-DKO mice (Suppl. Fig. 4, Suppl. Data 2) by an unbiased, data-independent acquisition (DIA) proteomic approach.

**Figure 4.**
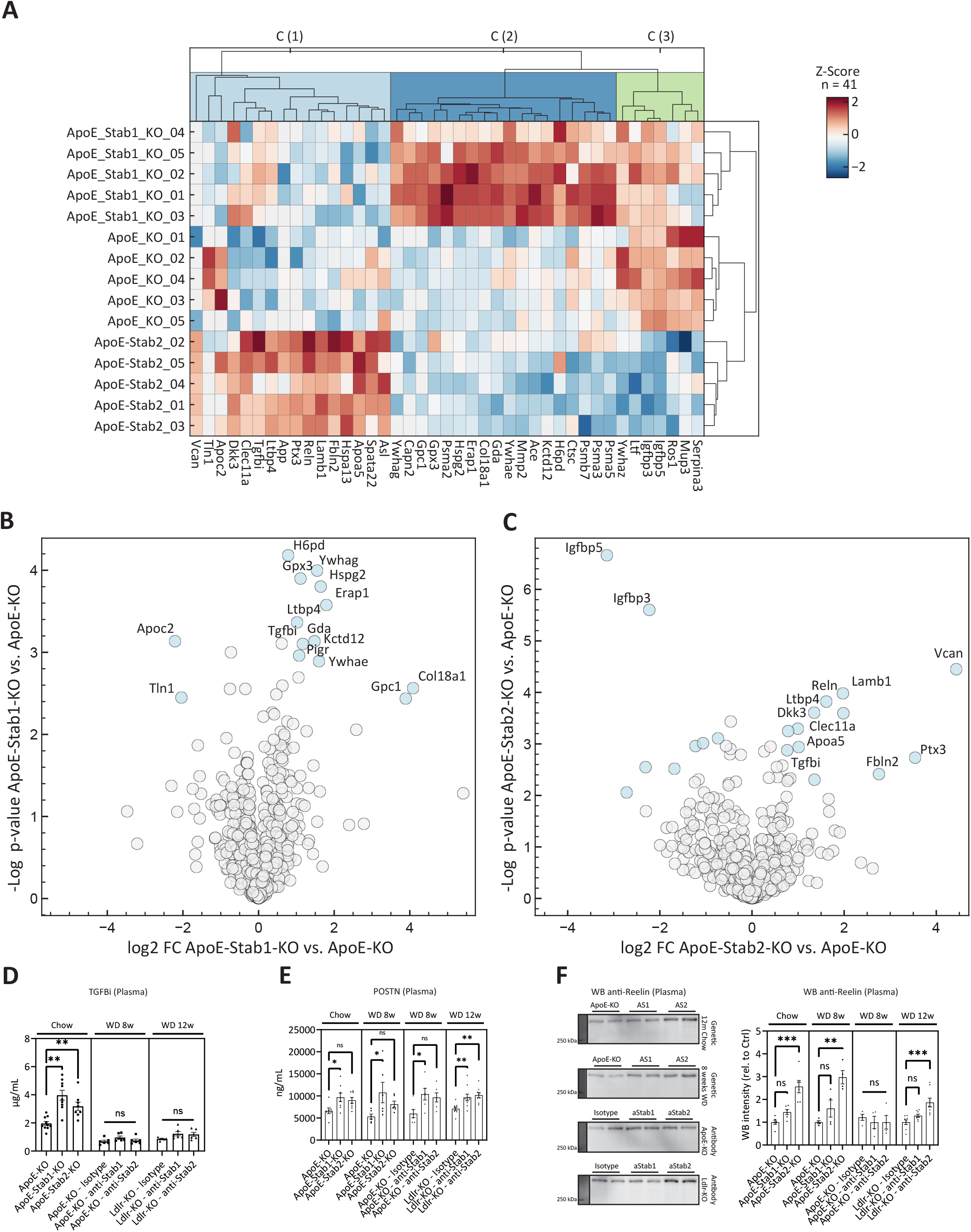
Proteomic plasma alterations in ApoE-Stabilin-KO mice **(A)** Clusters of 41 differential plasma proteins in ApoE-KO, ApoE-Stab1-KO and ApoE-Stab2-KO mice (n=5). C (1) (light blue) denotes Cluster 1 (strongest increase in ApoE-Stab2-KO), C (2) (dark blue) denotes Cluster 2 (strongest increase in ApoE-Stab1-KO), C (3) (light green) denotes Cluster 3 (strongest decrease in ApoE-Stab2-KO). **(B-C)** Volcano plots showing log2 fold changes of detected plasma proteins on the x-axis and the –log10 p-value (two-sided t-test) on the y-axis. **(B)** ApoE-KO vs. ApoE-Stab1-KO plasma. **C:** ApoE-KO vs. ApoE-Stab2-KO plasma. Significantly changed proteins (permutation-based FDR < 0.05) are highlighted in light-blue. **(D)** TGFBi (n≥4) was measured by ELISA in blood plasma of aged ApoE KO mice with Stabilin deficiency and anti-Stabilin-antibody treated mice. **(E)** Postn (n≥6) was measured by ELISA in blood plasma of aged ApoE KO mice with Stabilin deficiency and anti-Stabilin-antibody treated mice. **(F)** Reelin WB of plasma of chow-fed aged ApoE KO with genetic Stabilin deficiency as well as ApoE and LDLR KO treated with anti-Stab1 and anti-Stab2. Representative images (left panel) and quantifications by densitometry (right graph, n≥4). Error bars represent SEM. ns=not significant, *p<0.05, **p<0.01, ***p<0.001.

In ApoE-KO, ApoE-Stab1-KO and ApoE-Stab2-KO, 721 proteins were quantified, of which 539 (75%) were found in all samples. Altogether, 41 proteins were identified that showed a significant dysregulation among the three groups (Fig. 4A), of which 28 were found in all samples. Eight proteins (RELN, VCAN, PTX3, FBLN2, LAMB1, APOA5, CLEC11A, DKK3) were significantly more abundant in ApoE-Stab2-KO plasma samples, while 10 proteins (KCTD12, COL18A1, GPX3, YWHAG, YWHAE, H6PD HSPG2, ERAP1, GPC1, GDA) were significantly more abundant in ApoE-Stab1-KO compared to ApoE-KO. Among these, twelve proteins (H6PD, COL18A1, HSPG2, GPX3, GPC1, FBLN2, PTX3, LTBP4, CLEC11A, DKK3, TGFBi and Reln) showed at least a trend towards higher plasma abundance in both ApoE-Stab1 and ApoE-Stab2 KO mice compared to ApoE-KO suggesting that these proteins may represent common ligands of both Stabilins in this context and may be involved in systemic control of atherosclerosis by either Stabilin (Fig. 4A-C, Suppl. Data 1). The skewed proportion of increased versus decreased plasma proteins indicates that impaired scavenging by Stab1 and Stab2 predominantly causes a retention of plasma proteins in the circulation.

To further validate the influence of Stabilin deficiency on the plasma proteome, we compared WT, Stab1-KO, Stab2-KO and Stab-DKO plasma samples. 563 proteins were quantified, of which 416 (74%) were found in all samples. Altogether, 231 proteins were identified that showed a significant dysregulation among the four groups (Supp. Figure 4), 151 proteins showed a post-hoc significance in at least one group compared to ApoE-KO (Suppl. Data 2). Comparing significantly dysregulated proteins among the two independent proteomic experiments, we found 24 proteins overlapping in both comparisons. Postn, TGFBi and Reln, all known modulators of atherosclerosis, were strongly increased in Stab-DKO mice (Suppl. Fig 4B, Suppl. Data 2). Since Reln contains EGF-like-domains and TGFBi and Postn contain fasciclin domains, we hypothesized that they might be direct ligands of Stab1 and/or Stab2, as Stab1 and Stab2 also contain EGF-like and fasciclin domains.

### Circulating levels of Periostin (Postn), Reelin (Reln) and Transforming growth factor-β-induced (TGFBi) are controlled by Stab1 and Stab2

Among the most strongly altered proteins in the plasma proteome analyses, we identified Postn, TGFBi and Reln as candidate ligands potentially involved in the systemic control of atherosclerosis by the Stabilins. Notably, Postn, TGFBi and Reln are known to exert disease-modifying functions in different stages of atherosclerosis development. Therefore, plasma levels of Postn, TGFBi and Reln were assessed by ELISA and western blotting, respectively. TGFBi was significantly increased in the plasma of ApoE-Stab1-KO and ApoE-Stab2-KO in comparison to ApoE-KO (Fig. 4D) and also significantly increased in Stab-DKO mice in comparison to WT mice (Suppl. Fig. 4C). Postn was significantly more abundant in ApoE-Stab1-KO, ApoE-Stab2-KO and all anti-Stab1-treated models, while anti-Stab2 treatment only showed significant enhancement of Postn only in Ldlr-KO (Fig. 4E). Similar to TGFBi, only Stab-DKO had higher plasma levels of Postn in comparison to WT (Suppl. Fig. 4C). This indicates that TGFBi and Postn both represent common ligands of Stab1 and Stab2, which might compensate for each other in single knockouts. In certain settings of ApoE-KO and Ldlr-KO, deficiency or inhibition of one of the Stabilins was sufficient to increase circulating levels of Postn or TGFBi indicating that clearance of the molecules is also affected by these pro-atherogenic models. Reln, as assessed by Western blotting, was signifcantly elevated in the plasma of ApoE-Stab2-KO and anti-Stab2 treated Ldlr-KO mice (Fig. 4F) as well as in Stab-DKO and Stab2 KO mice (Suppl. Fig. 4D). These findings indicate that the regulation of Reln levels in the circulation depends more heavily on the presence of Stab2 than Stab1.

### Periostin (Postn), Reelin (Reln) and Transforming growth factor-β-induced (TGFBi) represent novel ligands that are routed to lysosomal degradation by stabilin-mediated scavenging

To assess whether Postn, TGFBi, and Reln directly bind to Stab1 and/or Stab2, GST-pulldown assays with different fragments of human Stab1 and human Stab2 as well as endocytosis assays with transfected CHO-cells expressing either red fluorescent protein (CHO-RFP), mouse Stab1 and RFP (CHO-RFP+mStab1) or mouse Stab2 and RFP (CHO-RFP+mStab2) were performed. Indeed, Postn and TGFBi both bound to the Fasciclin domains of Stab1 and Stab2 in GST-pulldown assays and both showed significant endocytosis with lysosomal uptake by CHO-RFP+mStab1 and CHO-RFP+mStab2, but not by CHO-RFP (Fig. 5A-D). Reln on the other hand showed strong lysosomal uptake by CHO-RFP+mStab2, but not by CHO-RFP+mStab1 and CHO-RFP, further confirming that Reln is primarily bound by Stab2, but not Stab1 (Fig. 5E). Antibody internalization assays confirmed the specific uptake of anti-Stab1 antibody to Stab1-expressing cells, anti-Stab2 antibody to Stab2-expressing cells and excluded non-specific scavenging of the respective antibodies (Fig. 5F). Overall, Postn, TGFBi and Reln represent novel circulating ligands controlled by Stabilin-mediated endocytosis and degradation in physiologic and pro-atherogenic conditions.

**Figure 5.**
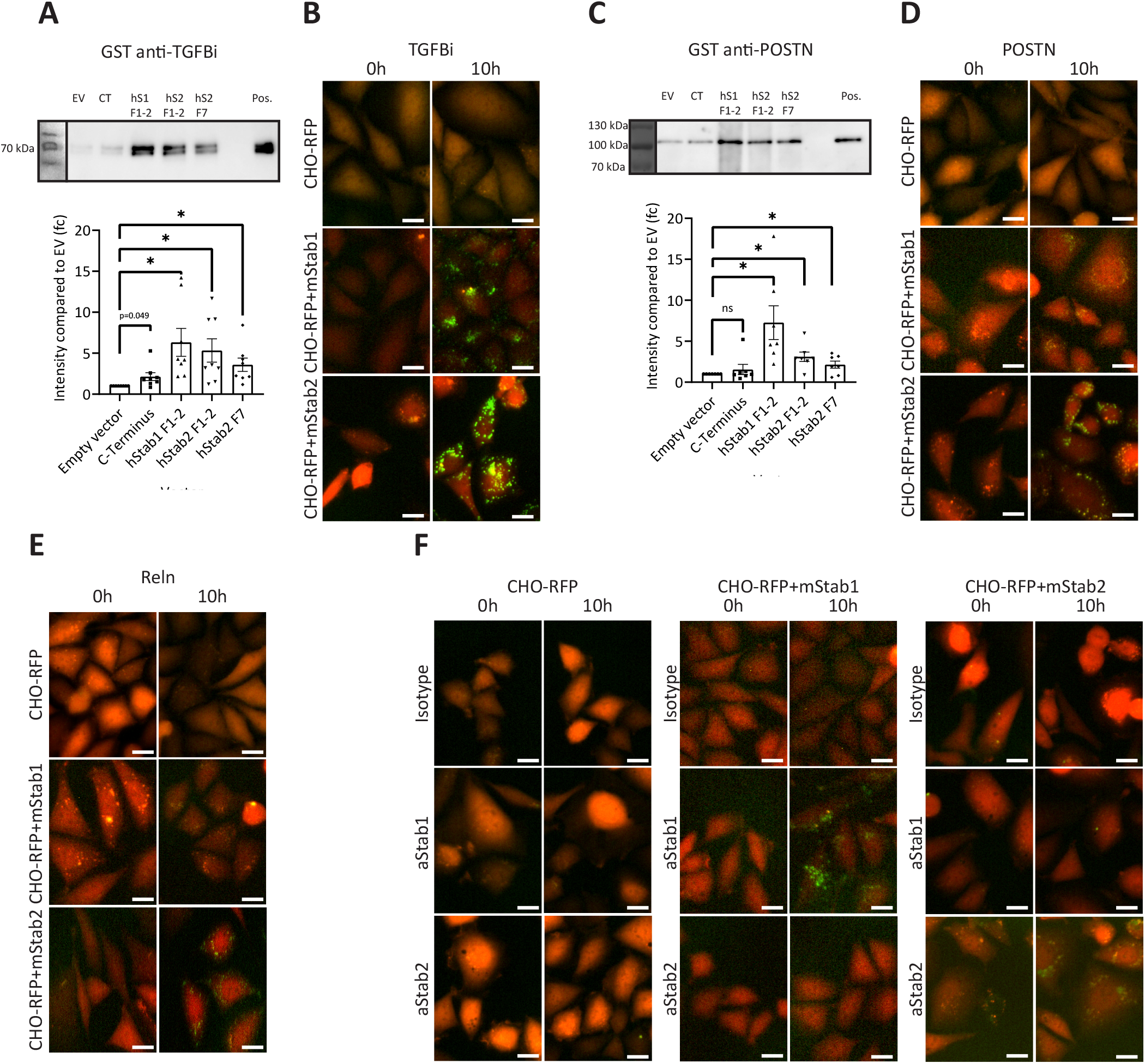
TGFBi, POSTN and Reln represent novel Stabilin ligands GST-pulldown with recombinant human TGFBi **(A)** or human POSTN **(C)** and GST-tagged fragments of human Stab1 and Stab2 (C-terminus, hStab1 F1-2, hSTab2 F1-2, hStab2 F7). Pos. indicates positive controls (recombinant protein). Representative WB images (upper panel) and quantifications by densitometry (lower panel, n≥6). Transfected CHO-cells expressing either RFP, mouse Stab1 and RFP or mouse Stab2 and RFP were treated with pHRodo green © labeled human TGFBi **(B)**, human POSTN **(D)**, human Reln **(E)**, isotype antibody, aStab1 antibody or aStab2 antibody **(F)**. Representative images are shown immediately after treatment (0h) and 10h after treatment. Green signal indicates activated pHRodo green © in cells. Error bars represent SEM. ns=not significant, *p<0.05, **p<0.01, ***p<0.001.

### Single cell RNA Sequencing reveals monocyte/macrophage suppression by down-regulation of Egr1 in patrolling as well as inflammatory monocytes

Since the slightly altered plasma lipid levels could not account for atheroprotection mediated by impairment of the scavenging functions of Stab1 and Stab2, we next considered modulation of systemic inflammation by plasma proteins as a potential mechanism for protection from atherosclerosis. When analyzed by flow cytometry, absolute and relative numbers of T-Cells, B-Cells and myeloid cells as well as lymphoid and myeloid cell subsets were not significantly altered (Fig. 6A, Suppl. Fig. 5A). Levels of inflammatory cytokines such as IFN-γ, IL-6, M-CSF, IL-1β and TIMP1 were assessed in the circulation of ApoE-KO, ApoE-Stab1-KO and ApoE-Stab2-KO mice as well as in the circulation of ApoE-KO and Ldlr-KO mice treated with anti-Stab1 and anti-Stab2 mAB. Notably, plasma levels of IFN-γ, IL-6, and M-CSF were significantly reduced in WD-fed ApoE-Stab1-KO and/or ApoE-Stab2-KO mice in comparison to WD-fed ApoE-KO mice, while plasma levels of IL-1β and TIMP1 were not altered (Suppl. Fig. 5B). Likewise, a trend to lower plasma levels of IFN-γ, IL-6, and M-CSF was also observed in ApoE-KO and less so Ldlr-KO mice treated with anti-Stab1 and anti-Stab2 mAB; these latter changes, however, were smaller and failed to reach statistical significance (Suppl. Fig. 5B).

**Figure 6.**
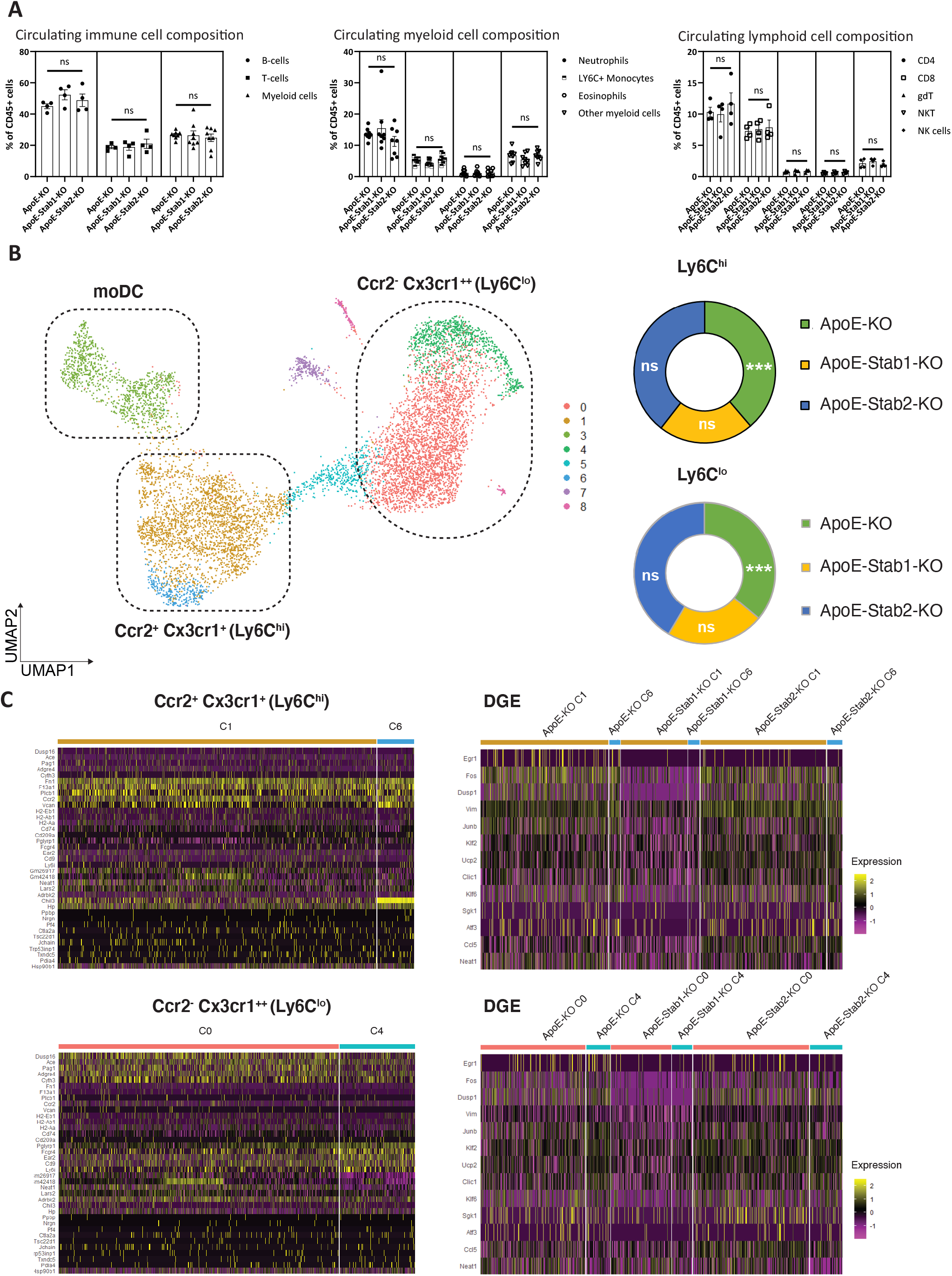
Circulating immune cells show differential gene expression in scRNAseq **(A)** Composition of circulating immune cells from the peripheral blood of 12 months old ApoE-KO, ApoE-Stab1-KO and ApoE-Stab2-KO on chow diet was assessed by flow cytometry. Frequencies of B-Cells, T-Cells and total Myeloid cells (left panel), Myeloid cell subpopulations (middle panel), and frequencies of Lymphoid cells (right panel) among gated immune (CD45+) cells are shown. **(B)** UMAP-Clustering of gated (live cells, CD45^+^ CD11b^+^ Ly6G^-^) myeloid cells without Cluster 2 (NK cells). Hypergeometric testing was performed for Ly6c-hi and Ly6c-lo monocytes to assess numeric composition adjusted to genotype. **(C)** Cluster-defining genes (left panels) and differentially expressed genes (right panels) of Ccr2^-^/Cx3cr1^++^/Ly6C^lo^ patrolling (Cluster 0 and Cluster 4) and Ccr2^+^/Cx3cr1^+^/Ly6C^hi^ inflammatory monocytes (Cluster 1 and Cluster 6) of ApoE-KO, ApoE-Stab1-KO and ApoE-Stab2-KO. Error bars represent SEM. ns=not significant, *p<0.05, **p<0.01, ***p<0.001.

As pro-atherogenic monocyte/macrophage activation is a hallmark of the development and progression of atherosclerosis ^35^, we reasoned that the beneficial effects of genetic or antibody-mediated targeting of Stab1 and Stab2 on plaque development might be driven by retention of atheroprotective factors in the circulation acting on monocytes/macrophages.

To this end, single-cell RNA-Sequencing (scRNAseq) of gated (live cells, CD45^+^ CD11b^+^ Ly6G^-^, Siglec-F^-^, CD19^-^, CD3^-^) myeloid cells of ApoE-KO, ApoE-Stab1-KO and ApoE-Stab2-KO was performed (Suppl. Figure 1 and 6). In these cells, no expression of Stab1 and Stab2 was found indicating that alterations in these cell populations cannot be related to cell-intrinsic deficiency of Stab1 or Stab2 (Suppl. Figure 6A). Unbiased clustering revealed 8 clusters in the sorted myeloid cells (Suppl. Fig. 6B-C, Suppl. Data 3). Excluding NK cells, the remaining seven clusters could mostly be summarized into Ccr2^-^/Cx3cr1^++^/Ly6C^lo^ (patrolling (Cluster 0 and Cluster 4)), Ccr2^+^/Cx3cr1^+^/Ly6C^hi (^inflammatory) monocytes (Cluster 1 and Cluster 6) and MoDC (Cluster 3) (Fig. 6B). Hypergeometric testing of patrolling and inflammatory monocytes showed a significant overrepresentation of both populations in ApoE-KO (Fig. 6B, Suppl. Data 4). When analyzing differential gene expression in Clusters among ApoE-KO, ApoE-Stab1-KO and ApoE-Stab2-KO genotypes (Suppl. Data 5), the most prominent alterations were seen in patrolling and inflammatory monocytes (Fig. 6C). Notably, pro-atherogenic Egr1 was the only gene commonly and significantly downregulated in both patrolling and inflammatory monocytes in ApoE-Stab1-KO and ApoE-Stab2-KO vs. ApoE-KO. Common Egr1 downregulation in monocytes in both ApoE-Stab1-KO and ApoE-Stab2-KO may thus provide a common mechanism reducing pro-atherogenic activation of monocytes (Supp. Figure 6D, Suppl. Data 6).

To validate the reduction of Egr1 expression on protein level in Cd11b^+^ cells, we performed Cytospins from peripheral blood of ApoE-KO, ApoE-Stab1-KO and ApoE-Stab2-KO mice, which were stained with anti-Cd11b and anti-Egr1 mAB. Here, a significant downregulation of Egr1 in Cd11b^+^ cells in ApoE-Stab1-KO and ApoE-Stab2-KO compared to Cd11b^+^ cells from ApoE-KO was confirmed (Fig. 7A-B).

**Figure 7.**
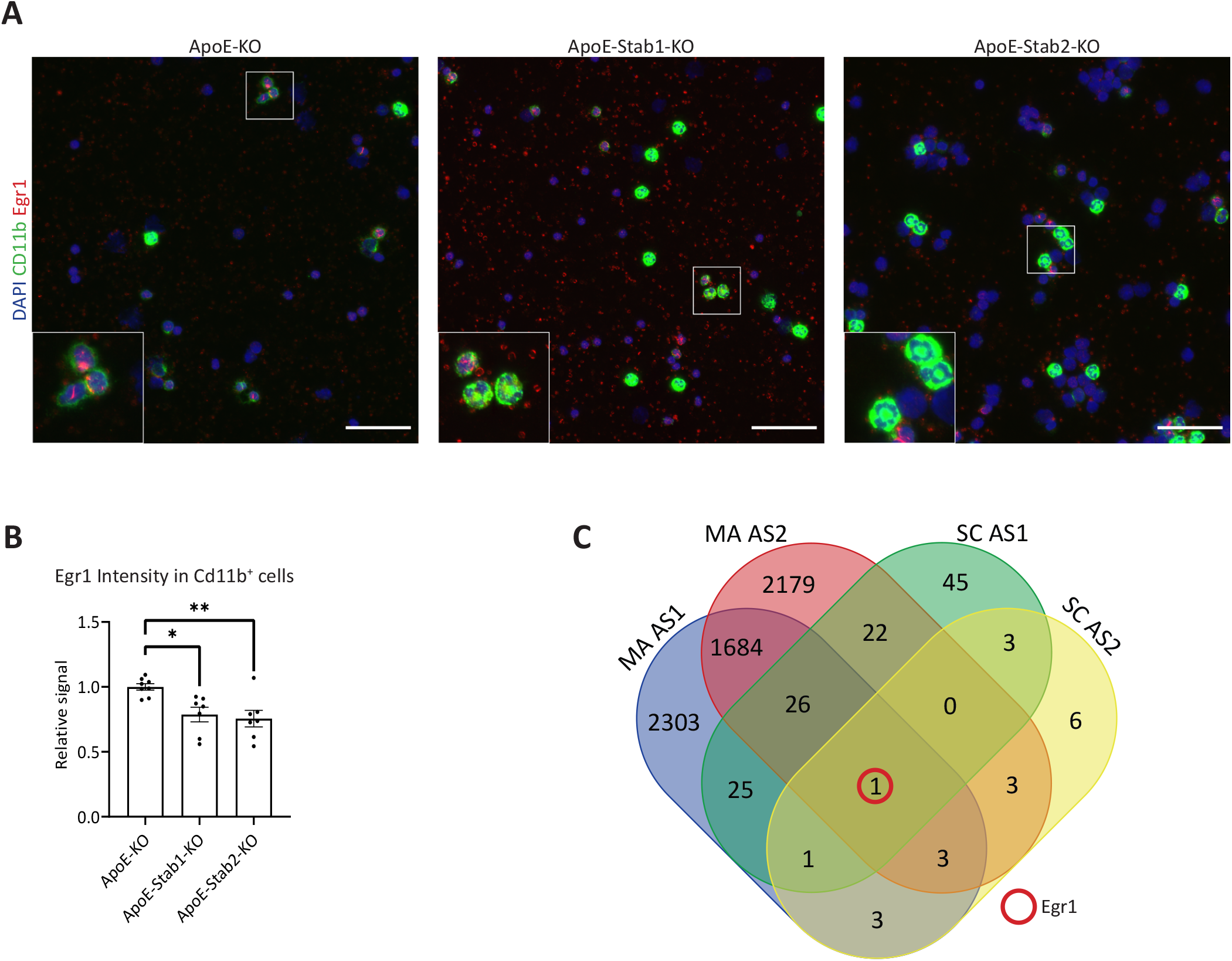
Egr1 is downregulated in monocytes *in vivo* and by blood plasma from Stabilin-deficient animals *in-vitro* **(A)** Representative immunofluorescent images of cytospins of white blood cells from ApoE-KO, ApoE-Stab1-KO and ApoE-Stab2-KO mice stained with DAPI (white), Cd11b (green) and Egr1 (red). Insets show Egr1 expression in Cd11b^+^ cells in different genotypes. **(B)** Comparative quantification of relative Egr1 immunofluorescence intensity in CD11b^+^ cells. n≥7 for both **(A)** and **(B)**. Scale bar (white) = 50 μm. **(C)** Venn-Diagram showing differentially expressed genes (adjusted p<0.05) among ApoE-Stab1-KO (MA AS1) and ApoE-Stab2-KO (MA AS2) plasma treated BMDM compared to APO-E KO plasma treated BMDM as well as differentially expressed genes (adjusted p<0.05) from scRNASeq in both Ly6c-hi (Cluster 1 and Cluster 6) and Ly6c-lo (Cluster 0 and Cluster 4) clusters in ApoE-Stab1 (*SC AS1*) and ApoE-Stab2 (*SC AS2*) compared to ApoE KO. Red circle indicates the only common regulation in all comparisons which is the downregulation of Egr1. Error bars represent SEM. ns=not significant, *p<0.05, **p<0.01, ***p<0.001.

To investigate how plasma protein changes may be linked to lower Egr1 expression *in-vivo*, bone-marrow-derived monocytes/macrophages (BMDM) isolated from WT mice were treated with recombinant TGFBi, Postn, Reln or a combination of the three ligands *in-vitro*. Treatment with these proteins induced no significant changes in Egr1 expression (Suppl. Fig. 7).

As a plethora of plasma proteins and cytokines were shown to be dysregulated, we reasoned that the more complex plasma protein changes, as indicated by proteomic analyses, are required to recapitulate the mechanisms mediating Egr1 downregulation *in-vivo*. Therefore, BMDM isolated from WT mice were exposed to blood plasma of WD-fed WT, ApoE-KO, ApoE-Stab1-KO and ApoE-Stab2-KO mice for 48 h. Global microarray mRNA gene expression analysis revealed 7225 dysregulated genes in BMDM (Suppl. Fig. 8, Suppl. Data 7).

1714 genes were commonly dysregulated in BMDM stimulated with ApoE-Stab1-KO or ApoE-Stab2-KO vs. ApoE plasma (yellow circle, Suppl. Fig. 8A). The majority of these commonly regulated genes showed parallel upregulation or downregulation in both BMDM stimulated with ApoE-Stab1-KO plasma or ApoE-Stab2-KO plasma in comparison to ApoE-KO plasma (∼70%). Likewise, KEGG pathway analysis revealed that except for one all the pathways commonly altered in BMDM stimulated with plasma of WT, ApoE-Stab1-KO and ApoE-Stab2-KO mice as compared to BMDM stimulated with plasma of ApoE-KO mice showed alterations of gene expression in the same direction as shown by NES (Suppl. Fig. 8B). Notably, exposure of mouse aortic endothelial cells (MAOEC) to these plasma samples did not similarly alter Egr1 levels indicating cell-specific effects occurring in BMDM, but not MAOEC (Suppl. Fig. 8C). Notably, combined analysis of differential gene expression data showed that Egr1 was the only gene commonly downregulated in Ly6c^hi^ and Ly6c^lo^ myeloid cells *in-vivo* and by ApoE-Stab1-KO and ApoE-Stab2-KO plasma treatment in BMDM *in-vitro* (Figure 7C, Supp Data 7). Reversion of Egr1-dependent pro-atherogenic monocyte activation via a plasma proteome switch thus likely explains protection from atherosclerosis through targeting of endothelial SR Stab1 and Stab2 *in-vivo*.

## Discussion

We here show that functional impairment of one of the Stabilins provides protection from aortic plaque progression in pro-atherogenic murine disease models, despite the fact that complete deficiency of both Stabilins impairs adult physiological homeostasis and survival ^16, 36^. We have recently shown that transplantation of Stab1-KO bone marrow into Ldlr-KO mice does not quantitatively alter aortic plaque development, indicating that Stab1-deficiency in monocytes/macrophages alone is not sufficient to generate the effects on atherosclerosis observed in the experiments presented here.

As Stab2 expression was not found in the aortic plaques and as Stab1 and Stab2 deficiency caused similar phenotypes of atheroprotection, it can be assumed that Stab1 and Stab2 rather exert remote, but not local control of atherosclerosis by regulating levels of circulating immunomodulatory molecules, including Stabilin ligands. This notion is also supported by additional findings. Firstly, scRNAseq of circulating myeloid cells did not show intrinsic expression of Stab1 or Stab2 despite common transcriptional alterations indicating reduced monocyte activation in patrolling and inflammatory monocytes in ApoE-Stab1-KO and ApoE-Stab2-KO including downregulation of Egr1. Secondly, ApoE-KO plasma-mediated changes in gene expression of BMDM are reverted in BMDM treated with plasma of ApoE-Stab1-KO and ApoE-Stab2-KO mice, including Egr1 repression. Egr1 has been shown to control proinflammatory and procoagulatory genes in monocytes and its deficiency has been reported to result in reduced atherosclerosis in Ldlr- and ApoE-KO mice^37, 38^. Common downregulation of Egr1 in monocytes/macrophages by plasma proteins controlled by Stabilin-mediated clearance is therefore a likely mechanism contributing to the atheroprotective effects observed. Overall, these findings demonstrate the striking functional significance of the Stabilin-controlled plasma proteome for atherosclerosis development.

Comprehensive proteomic analyses identified several new candidate ligands of Stab1 and Stab2 that may commonly mediate these effects by increased abundance in the circulation. Structural characteristics and similarities indicate that indeed several of these candidates may represent direct ligands. In this regard, Postn, TGFBi and Reln were all confirmed as direct ligands by us. All three have been shown to alter the outcome of atherosclerosis. TGFBi is one of four proteins in mammals with Fasciclin domains, the others being Postn, Stab1 and Stab2 ^39^. While Stab1 and Stab2 are transmembrane proteins, TGFBi and Postn are matrix proteins involved in tissue remodeling. Our data show that Stab1 and Stab2 represent the main scavenging system controlling levels of TGFBi and Postn in the circulation involving homotypic interactions of Fasciclin domains. Regarding the functions of TGFBi *in-vivo*, it is known that TGFBi mutations are found in corneal dystrophy ^40, 41^, and TGFBi is implicated in cancer progression ^42-44^. Elevated plasma levels of TGFBi are found in liver cirrhosis and non-alcoholic fatty liver disease ^45^. Surprisingly, TGFBi-null mice did not show gross corneal abnormalities, but lung alterations ^46, 47^. Although TGFBi was not studied in murine models of atherosclerosis, TGFBi expression is present in smooth muscle cells and macrophages in human atherosclerotic plaques ^48^. In contrast to TGFBi, Postn has been reported to be directly involved in the pathogenesis of atherosclerosis. Genetic deficiency of Postn protects Ldlr-KO mice from atherosclerosis ^49^. Mechanistically, this was related to alterations in the migratory behavior of macrophages. Postn variants are associated with atherosclerotic lesions in young people ^50^, and increased Postn deposition is found in human atherosclerotic lesions ^51^. The role of circulating Postn for atherosclerosis has not been specifically investigated. Notably, Postn and TGFBi have both been shown to bind αvβ3, e.g. to recruit tumor-associated macrophages ^52, 53^. In addition, immobilized Postn and TGFBi promote myeloid cell migration through αMβ2 integrin adhesion ^54-56^, which is also expressed on resident macrophages as well as M2 macrophages ^57^. Therefore, high circulating levels of Postn and TGFBi, as found in our disease models, might interfere with αMβ2 and αvβ3 integrin-mediated adhesion and migration of pro-atherogenic inflammatory cells to sites of atherosclerosis.

In contrast to Postn and TGFBi, loss of Reln has been described to protect from atherosclerosis by reducing leukocyte-endothelial adhesion and lesional recruitment ^58^. The impact of increased circulating levels of Reln, however, has not yet been studied in models of atherosclerosis. On the one hand, increase Reln levels could have pro-atherogenic effects that may be overridden by the effects of atheroprotective factors in the complex composition of the plasma proteome in ApoE-Stab2-KO mice. Alternatively, increased Reln levels in ApoE-Stab2-KO mice might interfere with atherosclerosis progression due to receptor saturation, futile overactivation of downstream signaling or by complex formation in the circulation.

Opposing changes in HA levels were observed in ApoE-Stab1-KO and ApoE-Stab2-KO mice. HA levels were high in all models with Stab2 deficiency or inhibition, which is in line with previous findings ^9, 16^. While reduced HA levels have been described to increase atherosclerosis susceptibility in mice due to glycocalyx damage ^28^, cell-type-specific inhibition of certain HA synthetase isoforms have been reported to have a beneficial effect ^29^. Although HA has a highly context-specific role in atherosclerosis development ^59^, the opposite effects on HA levels in Stab1 vs. Stab2 deficiency exclude HA as single common mediator of Stab1/2-mediated atheroprotection.

Divergent ligand concentrations in ApoE-Stab1-KO, ApoE-Stab2-KO and ApoE-KO/Ldlr-KO treated with anti-Stabilin antibodies as compared to Stabilin-KO mice without an atherogenic genetic background may be explained by latent impairment of Stabilin-mediated clearance in pro-atherogenic mouse models. In this regard, TGFBi was significantly elevated in plasma of ApoE-Stab1-KO and ApoE-Stab2-KO and Postn was significantly increased in plasma of ApoE-Stab1-KO and showed a trend towards increased levels in ApoE-Stab2-KO, while TGFBi and Postn were not significantly altered in Stab1-KO and single Stab2-KO. Latent impairment of Stabilin-mediated clearance, which may be present due to disturbed lipoprotein turnover in ApoE-KO and Ldlr-KO, could make these models more prone to plasma molecule alterations through antibody-mediated Stabilin inhibition or single genetic Stabilin deficiency. As impairment of Stab1 or Stab2 in pro-atherogenic mouse models affects levels of numerous circulating mediators, the net result of decreased immune activation is likely mediated by complex interactions of various Stabilin ligands, other altered plasma proteins and cytokines. This notion is supported by the fact that Egr1 repression in BMDM *in-vitro* could only be induced by ApoE-Stab1-KO and ApoE-Stab2-KO plasma but not by exposure to Postn, TGFBi, Reln or a combination of the three ligands alone. This indicates that either direct effects of these three ligands on Egr1 expression in monocytes *in vivo* may require involvement of other circulating co-factors only present in ApoE-Stab1-KO and ApoE-Stab2 plasma or may primarily involve independent other mediators. Such co-factors and independent mediators may themselves either be directly controlled by Stabilin-mediated clearance or may involve intercalated circulating or non-circulating cells responding to the altered proteome. Involved mechanisms may include altered signaling activities by direct binding, complex formation, competitive inhibition, induction or repression of secondary mediators and others. In this regard, other circulating proteins besides Postn, TGFBi and Reln are likely involved in the disease pathogenesis including the identified candidates H6PD^60^, COL18A1, HSPG2^61^, GPX3^62^, GPC1^63^, FBLN2^64^, PTX3^65^, LTBP4^66^, CLEC11A and DKK3^67-69^, most of which have been implicated in atherosclerosis and cardiovascular diseases.

In conclusion, enhancement of the effects caused by targeting Stab1 or Stab2 in the context of a pro-atherosclerotic metabolic state may make Stab1 and Stab2 promising candidates for the development of future treatment modalities for atherosclerosis.

Although we did not observe major side effects during anti-Stabilin antibody therapy such as weight loss, liver fibrosis or kidney damage during treatment periods of up to 12 weeks, possible future therapeutic applications will require surveillance for potential side effects, which may especially involve fibrotic organ damage. Alterations of plasma proteins have long been assumed to mostly represent non-functional bystanders. This view is currently changing, as it has been shown, that the composition of young blood plasma can have beneficial therapeutic effects in aged mice in regard to neural cell activity and brain functions, while aged plasma induces opposite effects in young mice ^70-72^, leading to first clinical trials applying young plasma infusions for Alzheimer’s disease ^73^. SR-targeted therapies like anti-Stab1 and anti-Stab2 offer an alternative approach to alter the proteome of the circulating blood plasma for therapeutic purposes.

Blood tests are the most common diagnostic procedures in clinical medicine. Current practice involves selecting specific proteins and peptides, depending on the patient’s individual clinical features. However, this practice may change as recent research suggests that global proteomic blood plasma analysis and application of machine learning may allow simultaneous, comprehensive and individualized assessment of multiple organ functions, aging, health related behaviors, disease stages as well as risk prediction ^74, 75^. Alterations of circulating proteins are usually attributed to either cellular secretion or leakage as primary determinants. Proteomic analyses of the blood plasma of Stabilin-deficient mice provide proof-of-concept that specific SR can have a profound impact on the circulating plasma proteome to alter global immune responses as well as disease susceptibility in distant organs. Thus, development and interpretation of comprehensive plasma proteomic diagnostic approaches will require consideration of scavenging functions as potential key drivers of associated alterations. In this regard, it is highly intriguing that circulating levels of STAB1, STAB2 and their ligands exhibit a sex-specific difference only in aged people. Elderly men display significantly decreased levels of both Stabilins, while common protein ligands, i.e. GDF-15, Postn and TGFBi, are all increased in comparison to elderly women (https://twc-stanford.shinyapps.io/aging_plasma_proteome/). As combined Stab1 and Stab2 deficiency in mice is associated with shortened lifespan ^16^, a sex-specific decline of their clearance functions may also be involved in sex-specific differences of human aging.

In summary, SR mediated endothelial clearance is involved in various physiologic functions, as well as disease states in distant organs providing opportunities for targeted adjustments of the plasma proteome for therapeutic applications.

## Acknowledgements

We thank Jochen Weber, Hiltrud Schönhaber, Maria Muciek, Camela Jost, Stefanie Riester, Janina Ritz and Christof Dormann for excellent technical support. We thank Christel Weiß and Niklas Kehl for the support of the statistical analysis.

## Supplemental Materials

1. Expanded Materials & Methods containing Supplemental References ^16, 76-89^
2. Online Figures 1-8
3. Supplementary Data 1-7

