## Supplementary figures and images for "Targeting of scavenger receptors Stabilin-1 and Stabilin-2 ameliorates atherosclerosis by a plasma proteome switch mediating monocyte/macrophage suppression"

### Supplementary Figures

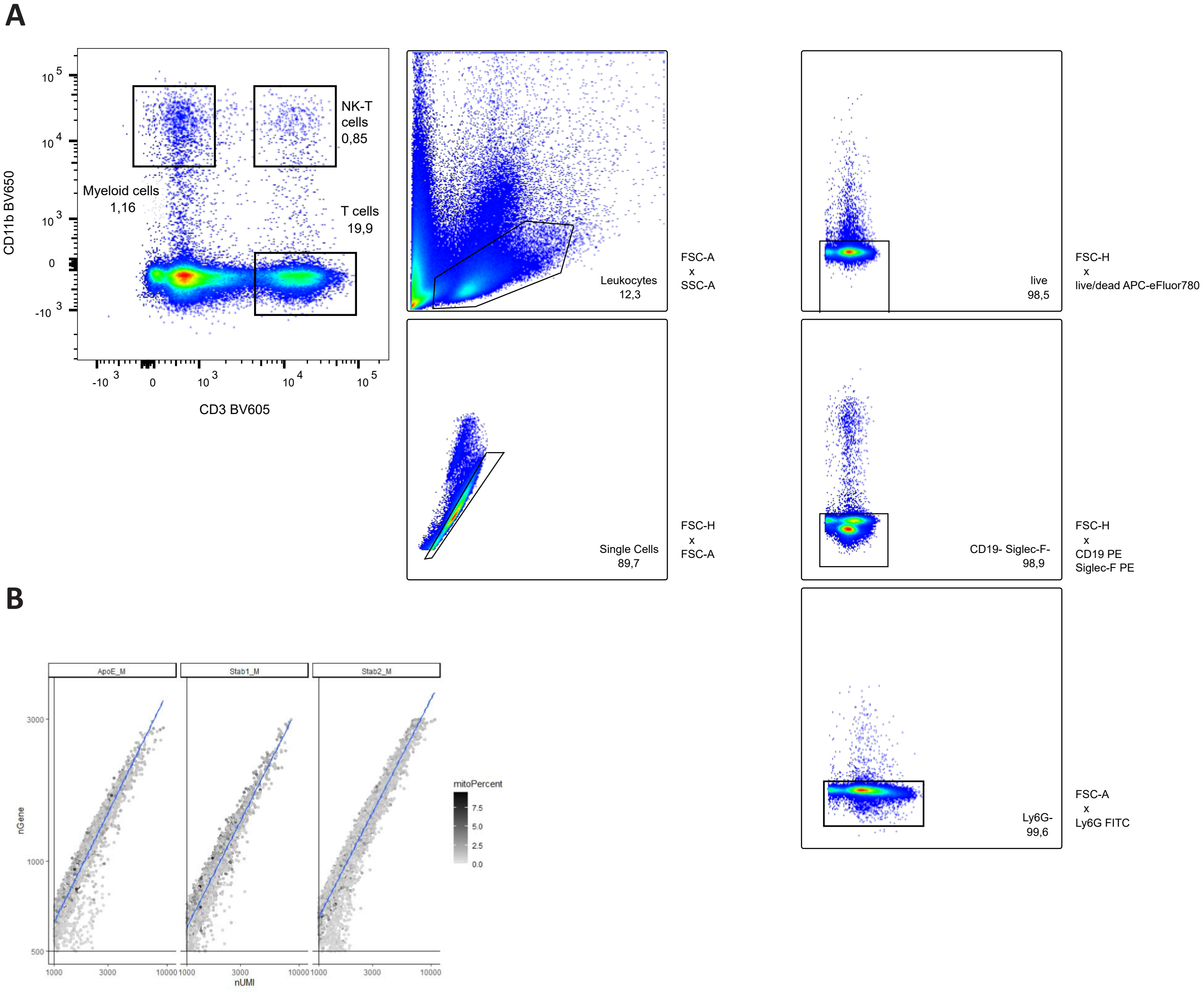

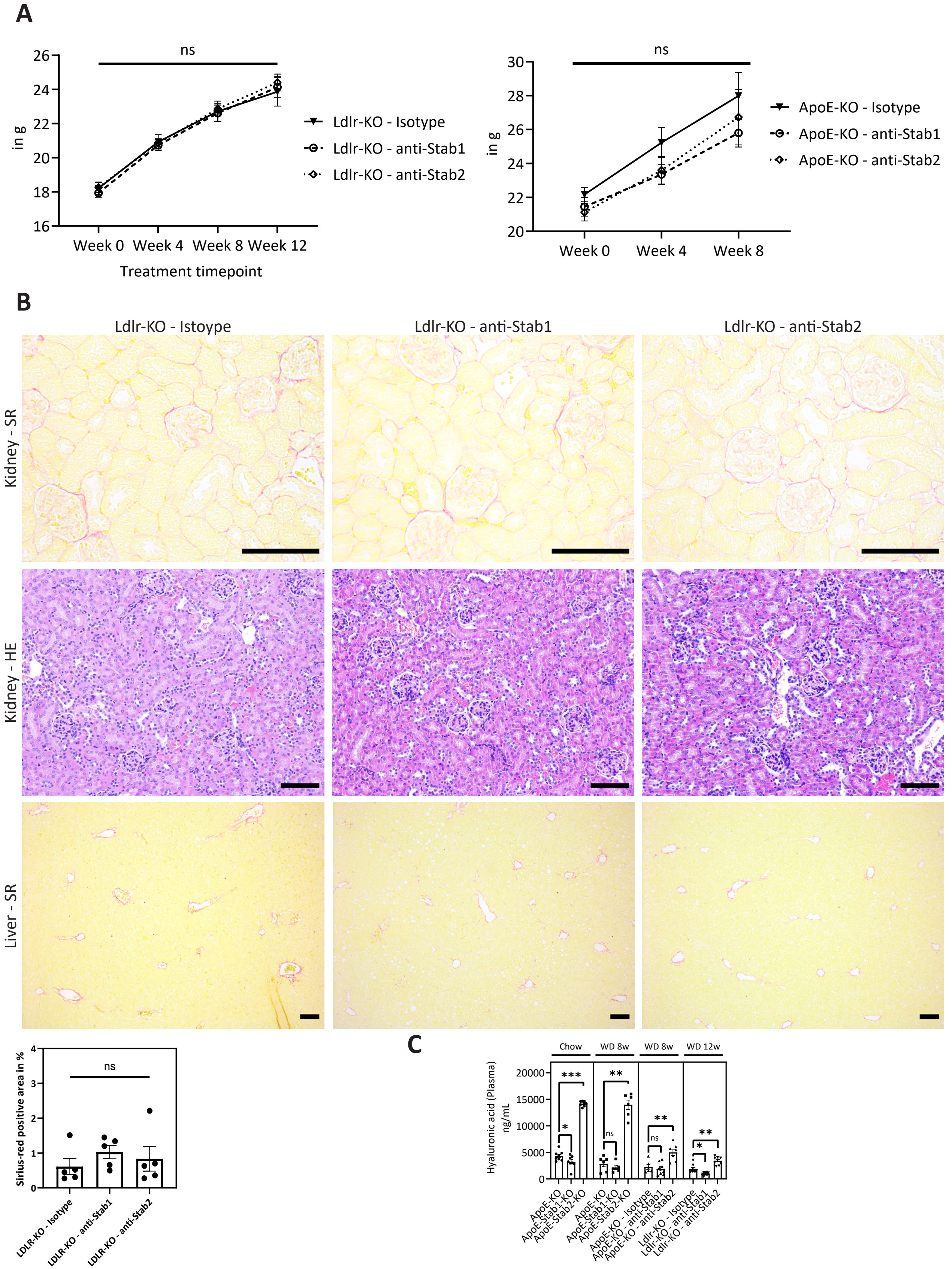

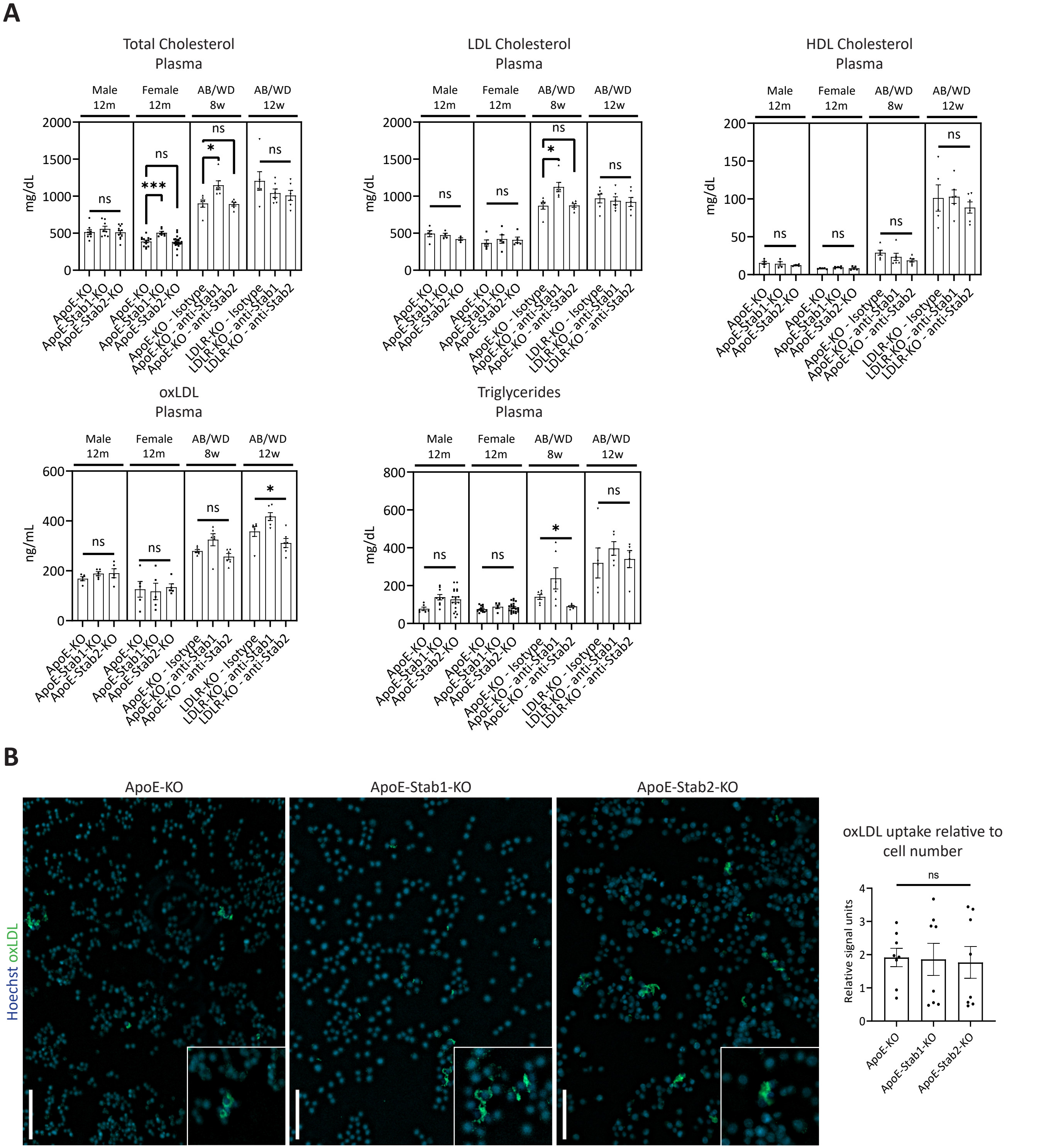

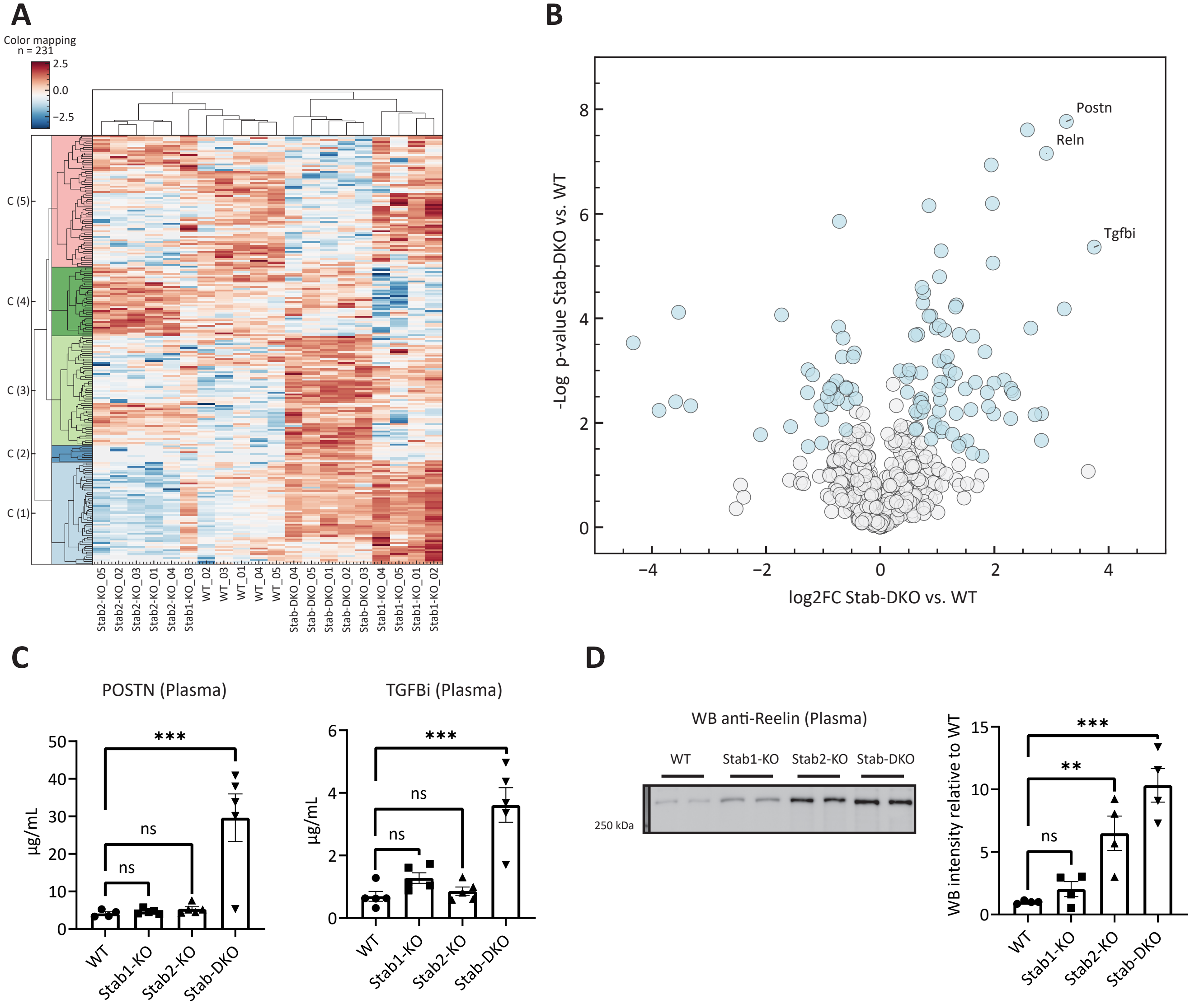

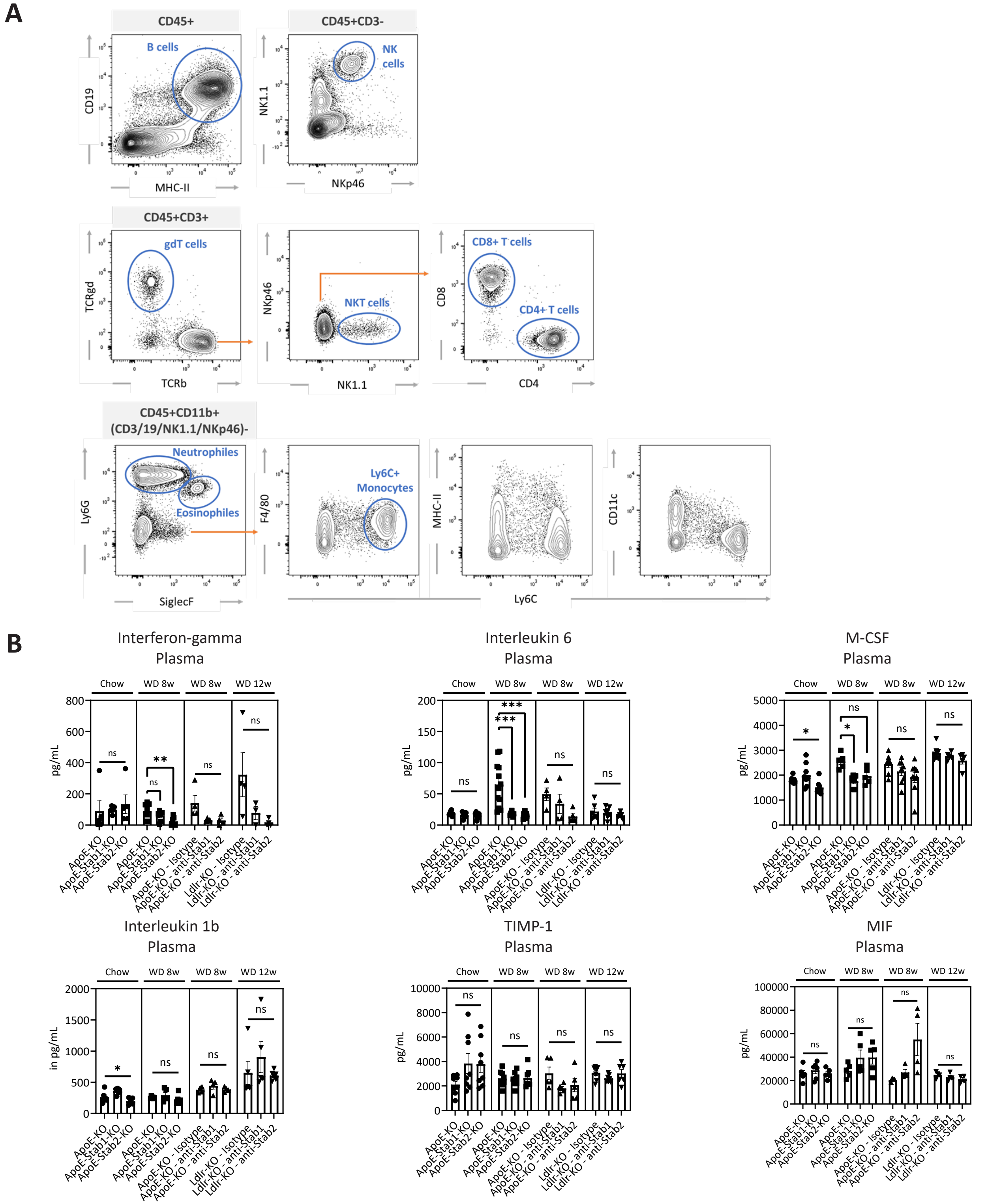

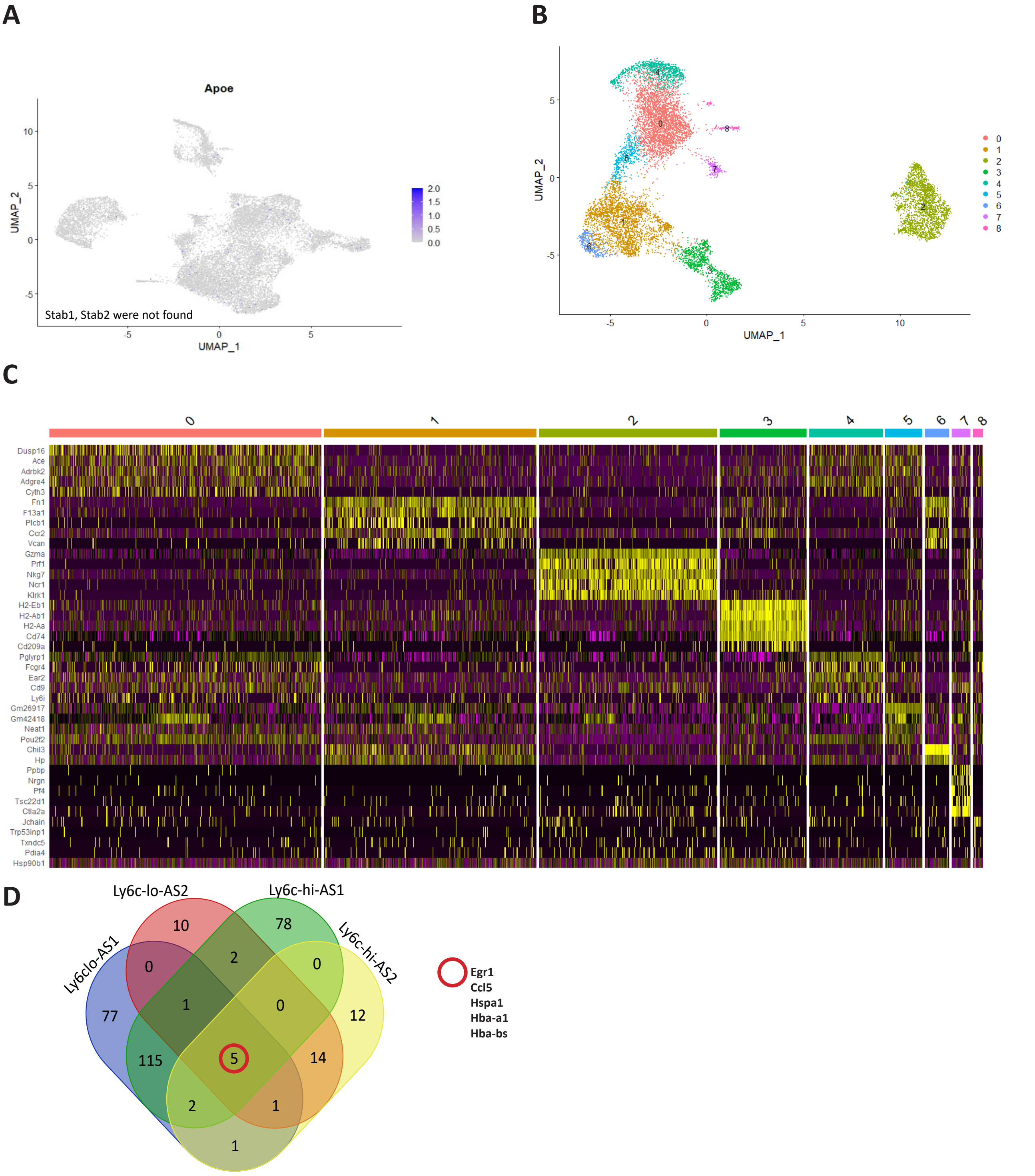

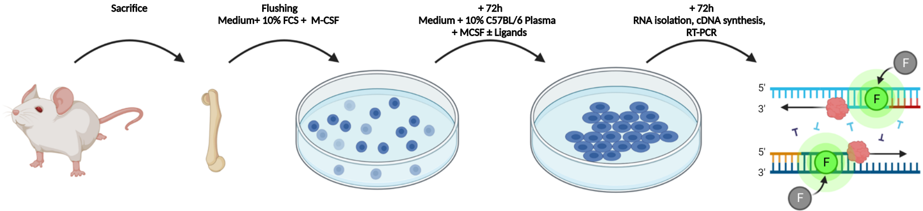

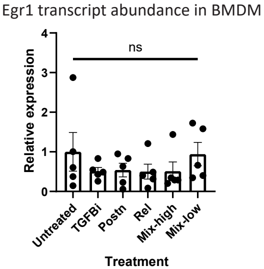

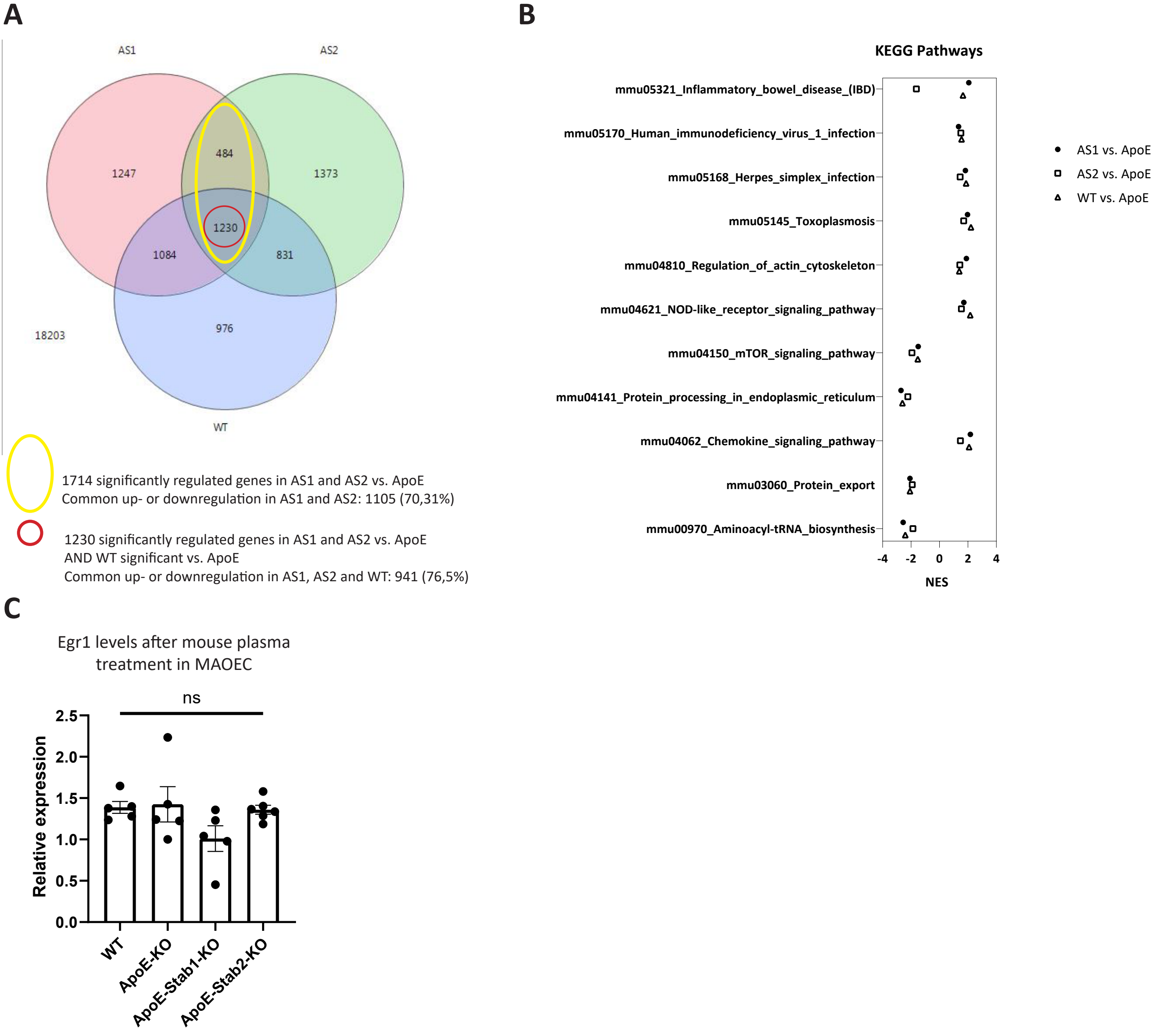
