## Supplementary material for "Targeting of scavenger receptors Stabilin-1 and Stabilin-2 ameliorates atherosclerosis by a plasma proteome switch mediating monocyte/macrophage suppression": Supp. Methods and Supplies. Fig. Legends

### Supplemental Materials

### Expanded Materials & Methods

*Animals.*

To investigate plaque formation in genetic models, ApoE-KO mice (B6.129P2-Apoe $^{tm1Unc}$ ,[76]) were crossed with Stab1-KO (B6.129S2-Stab1$^{tm1.1Cger}$ ,[16]) or Stab2-KO (B6.129S2-Stab2$^{tm1.1Cger}$, [16]) to generate transgenic mice with global deficiency for ApoE and Stab1 (ApoE-Stab1-KO) or ApoE and Stab2 (ApoE-Stab2-KO), respectively. To investigate plaque formation in antibody therapy models, ApoE-KO mice and Ldlr-KO mice (B6.129S7-Ldlr $^{tm1Her}$ [77]) were used. All mice were bred on a C57BL/6J background in the animal facilities of the Centre for medical research, Medical Faculty Mannheim. Ldlr-KO mice were purchased from Jackson at 8 weeks of age.

*Diets, anti-Stabilin antibody generation and anti-Stabilin-antibody treatment.*

To investigate spontaneous plaque formation, ApoE-KO, ApoE-Stab1-KO and ApoE-Stab2-KO mice were held on a regular chow diet (ssniff®R/M-H autoclavable, V1534-000, Ssniff, Germany) for twelve months. Western diet (Western Type Diet; TD88137; cat. no. DP110-03 ; SSNIFF) was fed to ApoE-KO , ApoE-Stab1-KO and ApoE-Stab2-KO aged 12 weeks for 8 weeks to induce plaque formation through dyslipidemia.

To generate monoclonal antibodies (mAB) against the extracellular N-terminal (NT) portion of Stabilin-1 and Stabilin-2, the corresponding mouse cNDAs encoding the first 711 (NT-Stabilin-1; NM_138672.2) or 730 amino acids (NT-Stabilin-2; NM_138673. 3), were PCR amplified and cloned by fusion with a murine Fc protein expression vector (PCR primers Stabilin-1: mSTB1_Bgl_Spe_fw 5' GAA GAT CTA CTA GTG TCT CAG ACA GC 3' and mSTB1_NotI_rv 5' ATG CGG CCG CAC AGC CTT TCT TCA 3'; pcr primers Stabilin-2: mSTB2_BamHI_Spe_fw 5' GAG GAT CCA CTA GTT GAA ATC GTG 3' and mSTB2_NotI_rv 5' TAG GCG GCC GCC ACA GTC ACA TCA C 3'). The CDNA3.1 IgG2b expression vector was kindly provided by S. Gordon [78]. Stable HEK293-stabilin-secreting clones transfected with lipofectiamine were selected with G418-

supplemented RPMI+5% FCS medium. High-purity NT-mStabilin-1-murineFc and NT-mStabilin-2-murineFc were recovered and concentrated from cell culture supernatants by protein A Sepharose FPLC chromatography and rebuffered in PBS. Subsequently, Stabilin-1 knockout and Stabilin-2 knockout mice (on a mixed 129;B6N genomic background) were immunized with NT-Stabilin-1 and NT-Stabilin-2, respectively, according to a standard routine immunization regimen for ten weeks (3x 100 µg/100 PBS mixed with incomplete Freunds adjuvant every 15 days). A final boost (without Freunds adjuvant) was placed 18 days after the third immunization, and spleen cells were harvested three days after that and PEG4000 fused with P3X63Ag8,653 cells. The fused hybridoma cells were cultured at a dilution of one cell per well in 96-well plates using HAT medium selection. Appropriate clones were screened with immunohistochemistry staining of liver tissue of RAG2 knockout mice using crude hybridoma supernatant. Clone 1.26 was selected as anti-mouse Stabilin-1 clone 1.26. Monoclonal antibody 1.26 is cross-reactive with rat tissue but not with human tissue. Anti-mouse Stabilin-2 monoclonal antibody clone 3.1 was prepared from immunized stabilin-2 knockout mice using the same protocols. The selected clone 3.1 is cross-reactive with rat and human Stabilin-2.

To evaluate the effects of anti-Stabilin antibodies on plaque formation, ApoE-KO and Ldlr-KO mice were injected twice weekly with 6 µg antibody per gram body weight with anti-Stabilin 1 (mSTB1 #1.26, InVivo BioTech Services GmbH cat. No. #AK726), anti-Stabilin 2 (mSTB2 #3.1, InVivo BioTech Services GmbH cat. No. #AK2810) or isotype IgG1 control antibody (Bioxcell; cat. no. BE0083) for the duration of Western diet feeding. ApoE-KO animals were injected and fed for 8 weeks at 12 weeks of age, Ldlr-KO animals for 12 weeks, starting at 8 weeks of age.

*RNA in situ hybridization.*

PFA-fixed liver tissue was sectioned at 4 microns. A modified non-isotopic in-situ hybridization protocol was carried out using the RNAscope 2.5 HD Red kit (Advanced Cell Diagnostics) following the manufacturer's recommended protocol with specific probes against the positive control mouse Ppib (Cyclophilin B) gene and mouse Stab1 or Stab2. Sections were ultimately stained with DAB and counterstained with hematoxylin.

*Plasma lipids.*

Mice were anesthetized, blood was collected by retrobulbar punction in microvette lithum heparin tubes (Sarstedt; cat. no. 201345). Plasma was separated (centrifugation at 2000 g for 5 minutes at 20 °C in Microvette 500 LH; Sarstedt) and analyzed for triglycerides and total cholesterol (cobas c 311 Analyser; Roche Diagnostics). Calibration and independent controls were used as recommended by the manufacturer. HDL and LDL were measured according to the manufacturers instructions by using a HDL and LDL/VLDL Cholesterol Assay Kit (STA-391; Cell Biolabs).

*Plasma Protein Digestion.*

Plasma samples were lysed in 4% SDS and reduced (TCEP, 10 mM) and alkylated (CAA, 20 mM) for 60 min at 45°C and digested using the SP3 digestion technique [79] using a one to one mixture of two different magnet beads (Sera-Mag(TM) Magnetic Carboxylate Modified Particles (Hydrophobic) # GE44152105050250, Sera-Mag(TM) Magnetic Carboxylate Modified Particles (Hydrophylic) # GE24152105050250). 3 µL of washed bead mixture was added to 20 µg of protein lysate and acetonitrile was added to a final concentration of 50%. The lysate was incubated for 8 min on a magnet and the liquid was removed. Proteins bound to the beads, were washed twice with 200 µL acetonitrile. Beads were dried for 3 minutes and 10 µL of a digestion solution containing 0.1 µg Trypsin (Sigma Aldrich) and 0.1 µg Lys-C (Wako) in 20 mM HEPES pH=8.5 was added for overnight incubation at 37°C.

Peptides were cleaned-up by 2 wash steps using 200 µL acetonitrile. Beads were incubated in 2% freshly prepared DMSO (Sigma Aldrich) and dried to complete dryness in a speed vac concentrator (Eppendorf). Prior to LC-MS/MS measurements, peptides were re-suspended in 10 µL 2% acetonitrile and 2% formic acid.

*Liquid Chromatography and Mass Spectrometry.*

LC–MS/MS instrumentation consisted out of an Easy nLC-1200 (Thermo Fisher) coupled via a nanospray-ionization source to an Exploris 480 (Thermo Fisher) mass spectrometer. A binary buffer system consisting out of solvent A and B (buffer A: 0.1% formic acid and buffer B: 0.1% formic acid in 80% acetonitrile) was utilized for peptide separation.

The in-house packed column length was 40 cm (ID = 75 µm). The column was filled with PoroShell C18 2.7-µm (Agilent Technologies) beads and a column oven controlled the temperate at 50 °C (PRSO-V2, Sonation). The buffer B percentage was linearly raised from 5%

to 27% within 69 min and further increased to 65% within 10 min. Buffer B content was then increased to 95% within 6 min. The column was washed at 95% B for 10 min. The total method time was 95 min per samples but the acquisition stopped after 90 min. All samples were measured in random order. The mass spectrometer operated in data independent acquisition mode. MS1 spectra were acquired at a resolution of 120,000 and an AGC target of $1 \times 10^6$. In total, 48 DIA windows were acquired at an isolation m/z range of 15 Th and the isolation windows overlapped by 2 Th, covering a mass range from 320 to 1040 m/z. Resolution of MS2 spectra was set to 15,000 at 200 m/z using a maximal injection time of 22 ms and stepped normalized collision energies (NCE) of 26, 28, 30. The samples were measured using the FAIMS interface at compensation values of -50 using the following temperature settings to decrease the FAIMS resolution: Inner electrode temperature: 99.5°C and the outer electrode temperature was set to 85°C. The FAIMS gas was set to 0 L/min (off) for WT, Stab1-KO, Stab2-KO and Stab-DKO samples. For ApoE-KO, ApoE-Stab1-KO and ApoE-Stab2-KO mice samples, a gas flow of 4 L/min was applied from 1-5 min gradient time.

*Proteomics Data Analysis and data Processing.*

Raw files acquired using DIA (Data independent acquisition) mode experiments. We utilized DIA-NN version 1.8 [80]. First a library was generated using all acquired raw files using the Mus Musculus Uniport reference proteome fasta file. The options 'Fasta digest for library-free search / library generation' and 'Deep learning-based spectra, RTs and IMs prediction' were enabled. A single miss cleavage was allowed and N-term M excision and carbamidomethylation at cysteine residues were defined as fixed modification. 'Mass accuracy', 'MS1 accuracy', and 'Scan window' options were set to 0 (automatic interference). The neuronal network classifier was to 'Double-pass mode'. The Precursor m/z range was defined to 300 – 1200. Further settings were 'Protein interence' = Isoform IDs and the quantification strategy was set to 'Robust LC (high accuracy)'. The generated library containing 5216 potentially precursors was then used to re-analyze the raw files. Quantities matrices were exported and the protein group specific output was used in the downstream analysis. To identify significantly different proteins, a two-sided t-test and if applicable an 1W-ANOVA was applied using log2 transformed MaxLFQ intensities [81] calculated within the DIA-NN software. The FDR was controlled to 5% using a permutation-based approach in the Perseus software. Results were visualized using the Instant Clue [82] software.

The mass spectrometry proteomics data have been deposited to the ProteomeXchange Consortium via the PRIDE [83] partner repository with the dataset identifier PXD029342.

*In vitro Protein-Protein interaction.*

GST pull-down assays were performed according to the manufacturer protocol using the MagneGST® Protein purification System (Promega). Fragments of human Stabilin-1 corresponding to the amino acids 395 to 645 (Stabilin-1 F1-2), and human Stabilin-2 corresponding to amino acids 405 - 712 (Stabilin-2 F1-F2) and 2369 - 2472 (Stabilin-2 F7) were cloned into the EcoRI/XhoI sites of the pGEX4T1 vector (Invitrogen). GST-fused proteins were expressed in *E. coli* strain BL21-CodonPlus-RIL (Stratagene) and purified under nondenaturing conditions [76]. Recombinant proteins were used as followed: TGFBi (10569-H08H, Sino-biological); Periostin (3548-F2-050, R&D), Reelin (8546-MR-050, R&D).. Eluates of GST pull-downs containing soluble Stab1 or Stab2 binding partners were analyzed by Western blotting. For live cell imaging, proteins (TGFBi, POSTN, Reln, aStab1, aStab2 and Isotype) were labeled with the pHrodo™ iFL Green Microscale Protein Labeling Kit (P36015, ThermoFisher) according to the manufacturers instructions.

*Western blotting.*

Eluates of GST pull-downs or mouse plasma were analysed by SDS-Page and Immuno-blotting on PVDF membranes as by the manufacturer protocol (Trans-Blot Turbo Transfer System, Bio-Rad). Mouse plasma was diluted 1:5 in dH2O before electrophoretic protein separation. Incubation with primary antibodies: anti-TGFBi (R&D AF2935), anti-Reelin (AF3820, R&D), anti-Postn (ab14041, Abcam), anti-His (ab18184, Abcam) was performed at 4°C overnight at a 1:1000 dilution in 5% nonfat dry milk (Bio-Rad No. 1706404). Incubation of HRP conjugated secondary antibody (different species, Dianova) was performed for 1 hour at room temperature. Chemiluminescence (Millipore WBLUF0500) intensity detection was performed with the Azure® c400 imaging system (Azure ® Biosystems) and quantified using ImageJ 1.47v [84].

*Enzyme-linked immunosorbent assay (ELISA).*

ELISAs were performed as recommended by the manufacturer using mouse plasma. Signals were measured using a Tecan Infinite 200 Pro. The following ELISA Kits were used: Periostin/

OSF-2 (RnD; MOSF20), Hyaluronic Acid (RnD; DHYAL0), INF-γ (RnD; SMIF00); IL-6 (RnD; M6000B); MIF (LSBio; LS- F5436); M-CSF (RnD; MMC00); TIMP-1 (RnD; MTM100); IL-1β (RnD; cat. no. DY401-05), TGFBi (ThermoFisher, EMTGFBI), oxLDL (CloudClone, SEA527Mu).

*Live cell microscopy and protein internalization.*

CHO-cells were transfected with lentiviral vectors carrying red fluorescent protein (RFP, control), mStab1 + RFP or mStab2 + RFP using the ADR3 vector. 5000 cells (CHO-RFP, CHO-RFP+mStab1 or CHO-RFP+mStab2) per well were seeded in 96 well plates and allowed to adhere for 24h. Cells were then treated with labeled protein (1:1000) and imaged immediately afterwards in the Incucyte S3 System (Sartorius) using the 2019B Rev2 Software using a 20x objective using the green (Emission Wavelength: 524 nm; Passband: [504,544] nm, Excitation Wavelength: 460 nm; Passband: [440,480] nm) and red (Emission Wavelength: 635 nm; Passband: [625,705] nm, Excitation Wavelength: 585 nm; Passband: [565,605] nm) filter sets. Another imaging was performed at 10h after treatment.

*Peripheral blood mononuclear cells (PBMC) isolation and flow cytometry.*

Mice were anesthetized, blood was collected by retrobulbar punction in microvette lithum heparin tubes (Sarstedt; cat. no. 201345). Red blood cells were lysed using buffered ammonium chloride potassium phosphate solution (ACK buffer) for 5-10 min at room temperature. For flow cytometry, dead cells were labelled with ZombieAqua solution (Biolegend) according to manufacturers instructions, followed by treatment with anti-CD16/CD32-producing hybridoma supernatant (10%) to block Fc receptors, prior to surface epitope staining. Following surface staining, cells were washed and fixation and permeabilization was carried out using eBioscience™ FoxP3 Fixation/Permeabilization Kit (Invitrogen) according to manufacturer's instructions. Intracellular staining was performed in Permeablization Buffer. Cells were washed, resuspended in PBS and analyzed using BD LSRFortessa™ cytometer (BD Biosciences). Data were analyzed by FlowJo™ Software.

*Single-cell RNA-Sequencing of murine myeloid cells, transcript quantification and analysis*

Cells from N=16 murine whole-blood samples (n=4 ApoE-KO, n=4 ApoE-Stab1-KO and n=4 ApoE-Stab2-KO) were isolated by FACS using the marker panel outlined in Suppl. Figure 1. Samples were processed following the 10x genomics single-cell sequencing procedure for 3' scRNA-Sequencing. Murine samples were pooled and sequenced in multiplexed libraries with up to 40.000 cells each per lane on an Illumina NovaSeq sequencing system (paired-end multiplexing run) at a depth of ~130,000-200,000 reads per cell.

Libraries after quality control (nUMI (corresponding to nFeature_RNA)>= 1000; nGene (corresponding to nFeature_RNA) => 500; nGene <= 3000; mitoPercent < 10, log10GenesperUMI (defined as: log10(nFeatures_RNA) /log10(nCount_RNA)) > 0.8) comprising a total of 14,985 cells (ApoE KO: 5,529; Stab1 ApoE KO: 3,179; Stab2 ApoE KO: 6,277) were integrated and analyzed using Seurat v3.2.1 [85]. The counts table of the merged Seurat object was filtered to keep only genes that are expressed in 10 or more cells.

Data analysis was performed using the Seurat v3.2.1 [85] workflow. The counts tables were filtered for features expressed by at least 3 cells and cells with at least 500 detected features, corresponding to the arguments min.cells=3 and min.features=500 in the CreateSeuratObject function call. The datasets were merged, scaled and normalized using the SCTranform function [86] with the function set to return 3000 variable features and regress out the percentage of mitochondrial genes and ribosomal genes, corresponding to the arguments, variable.features.n=3,000, and vars.to.regress= c("mitoPercent", "riboPercent") .

First level Uniform Manifold Approximation and Projection and shared nearest neighbors graph construction were performed on the top 20 principal components. Cluster were identified with the resolution set to 0.6. Differential gene expression analysis was conducted using the FindAllMarkers function of Seurat v3 [85]. In a second step, we removed two RBC cluster (1258 cells; Hbb-a1, Hbb-a2, and Hbb-bs) and two low-quality clusters (mitochondrial gene expression, no canonical marker genes; 778 cells) for analysis. The cleaned Seurat object was then used for further analysis.

Second level Uniform Manifold Approximation and Projection and shared nearest neighbors graph construction were performed on the top 50 principal components. Clusters were identified with the resolution set to 0.4. Differential gene expression analysis and condition-wise comparison was conducted using the Seurat functions FindAllMarkers and FindMarkers of Seurat v3.2.1 according to the provided vignettes on: https://satijalab.org/seurat/vignettes [85]. For comparisons between 2 conditions the FindMarkers function was used. Features with an average log fold change of 0.25 and adjusted p-value of 0.05 were considered significant.

*Cluster enrichment analysis*

Enrichment analysis for a given condition in a cluster was conducted using a hypergeometric test implemented in R under the phyper function. This test considers the number of cells from condition x in a given cluster or group of clusters with respect to all cells from condition x in the dataset, all cells from condition y in the dataset and the number of cells in a given cluster or group of cluster. We used it to calculate the probability that number n or more cells from condition x could be found in a given cluster or group of cluster by chance. Statistical significance was assumed for probabilities of <0.05. Correction for multiple testing was achieved using the Benjamini-Hochberg method.

*Detailed statistical analysis.*

Statistical analyses, excluding scRNASeq and unbiased proteomics, were performed with JMP® 14 and SAS® 9.4M7 (SAS Institute Inc.). For group comparisons, the one-way ANOVA was used when the normality assumption was met using the Shapiro-Wilk test. We used the Brown-Forsythe test to check for equal variances. In case of unequal variances, the Welch ANOVA was used. For post-hoc analyses after significant ANOVA p-values, Dunnett's test was used. If the normality assumption was not met, we used a Kruskal-Wallis test with Steel with control as a post-hoc test. Reference groups for Dunnett's test or Steel with control were ApoE-KO, isotype-treated ApoE-KO, isotype-treated Ldlr-KO, Wildtype (WT) or Empty Vector as indicated by braces in the respective graph. Differences between data sets with $P < 0.05$ in ANOVA, Kruskal-Wallis test and post-hoc tests were considered statistically significant.

One-way ANOVA or Welch ANOVA with Dunnett's test was used in Figure 1A (male mice, unstained brightfield microscopy), Figure 1C, Figure 2, Figure 3A (except CD11b in Ldlr-KO mice), Figure 4D-F, Figure 6A (except Neutrophils), Figure 7, Supp. Figure 2C (all groups except Ldlr-KO mice), Supp. Figure 3A (Total Cholesterol, HDL cholesterol and LDL cholesterol in ApoE-KO mice treated with anti-Stabilin mAB and oxLDL in ApoE-KO and Ldlr-KO mice treated with anti-Stabilin mAB), Supp. Figure 4C-D, Supp. Figure 5 (except M-CSF in ApoE-KO, ApoE-Stab1KO, ApoE-Stab2-KO mice fed WD 8W and Interferon-gamma in chow-fed mice) and Supp. Figure 8B.

Kruskal-Wallis test with Steel with control test was used in Figure 1A (Oil-red O stained aortae of male and female mice; female mice, unstained brightfield microscopy), Figure 1B, Figure 3A (CD11b in Ldlr-KO mice), Figure 6A (Neutrophils), Supp. Figure 2B, Supp. Figure 2C (only Ldlr-KO mice), Supp. Figure 3A (only Total Cholesterol, HDL cholesterol and LDL cholesterol in

ApoE-KO mice treated with anti-Stabilin mAB and oxLDL in ApoE-KO and Ldlr-KO mice treated with anti-Stabilin mAB), Supp. Figure 3B, Supp. Figure 5 (only M-CSF in ApoE-KO, ApoE-Stab1KO, ApoE-Stab2-KO mice fed WD 8W and Interferon-gamma in chow-fed mice) and Supp. Figure 7.

One-sample t-test was used in Figure 5 to check for equal means compared to empty vector (EV=1). For weight curves, we used mixed model statistical testing (SAS MIXED procedure) to account for terms for week, group, and their interaction. Fixed variables: week, treatment (group), interaction week X treatment (group); random variable: Mouse number.

*Mouse aorta isolation.*

Mice were sacrificed by CO2-inhalation. The inferior vena cava was opened, PBS was flushed through the left ventricle to prevent blood clotting. The aorta was cut at the diaphragm, dissected up to the left ventricle and photographed immediately after the procedure with a DMIL stereomicroscope (Leica). Afterwards, the aorta was either opened longitudinally and stained with oil red O or fixed in 4% Formaldehyde overnight with subsequent paraffin embedding. After fixation, the aorta was sectioned in the beginning of the descending aorta after the aortic arch and in two sections of the descending aorta in thoracic (around 5th intercostal artery) and in the area of the aortic hiatus (diaphragm) for Elastica-van-Gieson based plaque quantification.

*Oil-red O staining.*

The longitudinally cut aorta was briefly washed in PBS, subsequently immersed in water for five minutes and transferred to 60% Isopropanol. After five minutes of incubation in Isopropanol, the Aorta was stained for 10 minutes in 0.3 g/L Oil red O-solution in 60% Isopropanol. After a short immersion in 60% Isopropanol after the staining procedure, the aorta was washed in water for 1 minute and photographed immediately after the procedure with a DMIL stereomicroscope (Leica). Aortic root sections were cut at 8 µm and stained with Oil-red O according to established protocols [87]

*Generation of bone marrow derived macrophages (BMDM).*

Bone marrow derived macrophages (BMDM) were generated from C57BL/6J mice as described previously [88]. In short, mouse tibia and femurs were dissected, filtered through a 70 µm Nylon cell strainer and washed once with PBS. After red blood cell lysis the cells were cultured in DMEM containing 10% FCS; 0.5% Penicillin; 0.5% Streptavidin and MCSF 30 ng/ml (Peprotech; cat. no. 315-02). At day four the initial medium was replaced. For protein stimulation experiments, the BMDM were further cultured with DMEM containing 10% mouse plasma and varying concentration of Reelin (0.5 µg/mL, Biotechne, 3820-MR/CF), TGFBi (1 µg/mL, Biotechne, 2559-BG), POSTN (1 µg/mL, Biotechne, 2955-F2) or a combination of these proteins (Mix hi: 0.5 µg/mL Reelin, 1 µg/mL TGFBI, 1 µg/mL POSTN, Mix-lo: 0.115 µg/mL Reelin, 0.23 µg/mL TGFBI, 0.23 µg/mL POSTN). No proteins were added in the control group. The RNA was isolated 72 hours after cultivation.

*Murine aortic endothelial cell cultivation and treatment*

Primary mouse aorta endothelial cells (MAOEC) (Pelo Biotech; cat. no. C57-6052) were grown to confluence in 6-well plates supplied with Endothelial Cell Medium (Pelo Biotech) containing 10% FCS. Subsequently, the Endothelial Cell Medium was supplemented with 10% mouse plasma as described below.

*Plasma stimulation experiments*

For plasma stimulation experiments, BMDM and MAOEC were further cultured with DMEM (BMDM) or Endothelial Cell Medium (MAOEC) containing 10% plasma of either WT, ApoE-KO, ApoE-Stab1-KO or ApoE-Stab2-KO mice after 8 weeks of Western Diet treatment. The RNA was isolated 48 hours after cultivation with the corresponding plasma.

*Foam cell formation assay*

RAW264.7 cells were cultured in RPMI 1640 Medium (glutamine supplemented as stock) with 10% FCS and 100 U of penicillin/ml, and 100 µg of streptomycin/ml. 1x $10^5$ cells per well in 500 µL medium were transferred to 24 well plates and incubated for 24 hours. Subsequently, RAW264.7 cells were incubated with 10% ApoE-KO, ApoE-KO-Stab1 or ApoE-Stab2-KO plasma, respectively. After 4 hours, DyLight™-488-labeled oxLDL was added to each well according to manufacturer's instructions (Cayman Chemical, cat. No. 601180) and incubated for 16 hours. After wahing steps, 500µL of Hoechst Dye was added, incubated for 15 minutes, followed by another washing step. Subsequently, pictures were acquired using an inverted cell culture microscope (Vert.A1, Zeiss) using the GFP and DAPI filters. Fluorescence at excitation 493 nm and emission 518 nm (DyLight™-488-labeled oxLDL) as well as 350 nm and 461 nm (Hoechst) was measured using a Tecan Infinite 200 Pro. GFP fluorescence intensity was normalized to Hoechst intensity to correct for cell densities.

*Cytospins*

Blood samples (200µL) were taken from the facial vein from aged (6-12 months old) ApoE-KO, ApoE-Stab1-KO and ApoE-Stab2-KO mice in EDTA tubes (Sarstedt, Microvette 200, 20.1288). Red blood cells were lysed using buffered ammonium chloride potassium phosphate solution (ACK buffer) for 5-10 min at room temperature. 4% of isolated cells were cytocentrifuged and frozen at -80°C.

*Immunofluorecence*

Aortic root cryosections (8 µm) and peripheral blood cells were fixed with phosphate-buffered 4 % paraformaldehyde (PFA) (0335, Carl Roth) for 10 min. Antibodies were diluted in Dako antibody diluent (S202230-2, Agilent Technologies)

For immunofluorescence staining, primary antibodies were incubated over night at 4 °C. After three washing steps with phosphate-buffered saline (PBS) (A0964.9050, VWR International, Radnor, PA, USA), fluorophore-conjugated secondary antibodies were incubated for one hour at room temperature. Sections were mounted with Dako fluorescence mounting medium (S302380-2, Agilent Technologies). Egr1 signal intensity was quantified in extrapolated $Cd11b^+$ cell area using ImageJ 1.47v [84].

*Microarray processing and statistical analysis*.

Gene expression profiling was performed using arrays Mogene-2.0-st from Affymetrix. cRNA synthesis and further biontinylated antisense cDNA was carried out according to the standard labelling protocol with the GeneChip® WT Plus Reagent Kit and the GeneChip® Hybridization, Wash and Stain Kit (both from Thermo Fisher Scientific). Hybridisation to arrays was performed on a GeneChip Hybridization oven 640, then dyed in the GeneChip Fluidics Station 450 and thereafter scanned with a GeneChip Scanner 3000. All of the equipment used was from the Affymetrix-Company (Affymetrix, High Wycombe, UK). The raw fluorescence intensity values were normalised applying quantile normalization and RMA background correction. Arrays were annotated with a custom CDF version 22 with ENTREZ based definitions. Differential gene expression was analysed with OneWay-ANOVA, using a commercial software package SAS JMP7 Genomics, version 6, from SAS (SAS Institute). A false positive rate of a = 0.05 with false discovery rate correction was taken as the level of significance.

To determine whether defined lists (or sets) of genes exhibit a statistically significant bias in their distribution, we performed a Gene Set Enrichment Analysis (GSEA) using the *fgsea* package (Sergushichev, 2016) for R (ssoftware R v3.4.0, R Core Team 2017). Pathways belonging to various cell functions such as cell cycle or apoptosis were obtained from public external databases (KEGG, http://www.genome.jp/kegg).

*RNA isolation.*

Bone marrow derived macrophages and murine aortic endothelial cells were washed with PBS and harvested in RLY Lysis Buffer (Bioline) The RNA isolation was performed with microRNA Isolation kit II (Bioline; cat. no. Bio-52075), according to the manufacturer's instructions.

*Reverse transcription and quantitative polymerase chain reaction (RT-qPCR)*

Reverse transcription (RT) into complementary DNA (cDNA) was conducted using Maxima Reverse Transcriptase (EP0752, Thermo Fisher Scientific) and Oligo(dT)18 primer (SO131, Thermo Fisher Scientific). Quantitative PCR (qPCR) was performed using innuMIX qPCR SyGreen Sensitive (845-AS-1310200, Analytik Jena, Jena, Germany) on a qTOWER 3 G touch thermal cycler (Analytik Jena). Primers for RT-qPCR were designed using NCBIs PrimerBLAST (https://www.ncbi.nlm.nih.gov/tools/primer-blast/). If possible, primer pairs were designed to span an exon-exon junction, thus being mRNA-specific. If this was not possible they needed to be separated by at least one intron. Primers were validated using no template controls, no RT controls, agarose gel electrophoresis and melt curve analysis. Amplification data were analyzed using qPCRsoft 4.0.8.0 (Analytik Jena). Normalized expression values were calculated using the Pfaffl method considering amplification efficiency values determined by standard curves. Reference gene PPIB was used as a reference gene based on its unaltered expression in Microarray and single cell analysis and previous publications [89].

*Histology.*

For hematoxylin & eosin (H&E), periodic acid-Schiff (PAS), Prussian blue and Sirius red staining, formalin-fixed, paraffin-embedded samples were processed according to standard protocols. Sirius red signal was quantified using channel separation and thresholding using ImageJ 1.47v [39].

**Supplemental Figure legends**

**Supplementary Figure 1.** Gating strategy for pre-sorting of single-cell RNA-Sequencing of murine myeloid cells and generation of GenePlots

**(A)** Exemplary gating strategy for myeloid cells isolated from murine whole-blood samples by fluorescence-activated cell sorting (FACS). Myeloid cells were defined as live cells, $CD45^+$ $CD11b^+$ $Ly6G^-$, Siglec-$F^-$, $CD19^-$, $CD3^-$. N=16 murine whole-blood samples (n=4 ApoE-KO, n=4 ApoE-Stab1-KO and n=4 ApoE-Stab2-KO). **(B)** GenePlot of nGene/nUMI for each multiplexed scRNA-Seq library (n=4 biological replicates of each genotype).

**Supplementary Figure 2.** anti-Stabilin-antibodies do not show obvious adverse effects after therapy for 12 weeks

**(A)** Body weight of antibody-treated mice (ApoE-KO, Ldlr-KO) was measured every four weeks during concomitant WD-feeding (n=10 for Ldlr-KO, n=6 for ApoE-KO). ns = no significance in comparison between groups using mixed model statistical testing to account for terms for week, group, and their interaction. Fixed variables: week, treatment (group), interaction week X treatment (group); random variable: Mouse number.

**(B)** Sirius-Red (SR) stained kidneys (upper panel), HE-stained kidneys (middle panel) and SR stained livers (lower left panel) of 5 months old Ldlr-KO mice on WD treated with Isotype, anti-Stab1 and anti-Stab2 for 12 weeks. Representative microscopic photographs (all n=5). SR stained areas as indicators of collagen content were quantified in hepatic microphotographs using ImageJ (right graph, n=5). Scale bar (black) = 100µm.

**(C)** HA levels were measured by ELISA of plasma of antibody treated ApoE and LDLR KO as well as ApoE KO with genetic Stabilin deficiency (lower right graph, n≥6).

Error bars represent SEM. ns=not significant, *p<0.05, **p<0.01, ***p<0.001.

**Supplementary Figure 3** Plasma lipids in mouse models and foam cell formation.

**(A)** Plasma lipid levels of in ApoE KO mice with genetic Stabilin deficiency and anti-Stab1 and anti-Stab2 treatment. Total Cholesterol (n≥6), LDL (n≥4), HDL (n≥4), oxLDL (n≥5) and Triglycerides (n≥5) were assessed.

**(B)** Representative microscopic photographs of RAW264.7 cells incubated with mouse plasma and DyLight™-488-labeled oxLDL (green) as well as Hoechst (blue). Quantification of oxLDL fluorescence relative to Hoechst fluorescence is shown on the right (n=8). Scale bar = 100 µm Error bars represent SEM. ns=not significant, *p<0.05, **p<0.01, ***p<0.001.

**Supplementary Figure 4.** Proteomic plasma alterations in Stabilin-KO mice

**(A)** Clusters of significantly altered (ANOVA, FDR < 5%) plasma proteins in WT, Stab1-KO, Stab2-KO and Stab2-DKO mice (all groups n=5, see also Suppl. Table 2). Cluster analysis revealed that the most prominent alterations in blood plasma protein composition were found in Stab-DKO mice, with proteins in clusters 1-4 strongly increased in this group. Additionally, proteins in cluster 4 showed preferential accumulation in the blood plasma of Stab2-KO mice, proteins in cluster 1 and 5 in Stab1-KO, respectively.

**(B)** Volcano plot showing log2 fold changes of differentially distributed plasma proteins in Stab-DKO vs. WT plasma on the x-axis and the -log p-value (two-sided t-test) on the y-axis (n=5). Significantly (permutation-based FDR < 0.05) changed proteins are highlighted. The plasma proteins most strongly upregulated in this comparison comprised Postn, Tgfbi and Reln.

**(C)** Postn (left graph) and TGFBi (right graph) were measured by ELISA in plasma of WT, Stab1-KO, Stab2-KO and Stab-DKO mice (n≥4).

**(D)** Western blot (WB) of plasma of WT, Stab1-KO, Stab2-KO and Stab-DKO with anti-Reelin antibody. Representative image (left graph) and quantification by densitometry (right graph, n=4).

Error bars represent SEM. ns=not significant, *p<0.05, **p<0.01, ***p<0.001.

**Supplementary Figure 5.** Gating strategy for FACS and Plasma Cytokines in different mouse models

**(A)** Indicated immune cell subsets were defined as shown, among pre-gated live CD45+ immune cells.

**(B)** INF-g (n≥3), IL-6 (n≥4), M-CSF (n≥5), IL-1b (n≥4), TIMP-1 (n≥5) and MIF (n≥4) as measured by ELISA of blood plasma of ApoE KO with genetic Stabilin deficiency as well as ApoE and LDLR KO treated with anti-Stab1 and anti-Stab2.
Error bars represent SEM. ns=not significant, *p<0.05, **p<0.01, ***p<0.001.

**Supplementary Figure 6.** Single-cell RNA-Sequencing of murine myeloid cells.
**(A)** DimPlot showing expression of ApoE in only very few of all sequenced cells. No transcripts of Stab1 or Stab2 were detected in the merged dataset.
**(B)** UMAP-Clustering of gated (live cells, CD45+ CD11b+ Ly6G-) myeloid cells containing all cells.
**(C)** Heatmap of cluster defining genes for all cells and cluster-defining genes.
**(D)** Venn-Diagram showing differentially expressed genes (adjusted p<0.05) among (adjusted p<0.05) from scRNASeq in Ly6c-hi (Cluster 1 and Cluster 6) and Ly6c-lo (Cluster 0 and Cluster 4) clusters in ApoE-Stab1 (*Ly6c-lo-AS1* and *Ly6-hi-AS1*) and ApoE-Stab2 (*Ly6c-lo-AS2* and *Ly6-hi-AS2*) compared to the respective groups in ApoE KO. Red circle indicates the only common regulation in all comparisons which is the downregulation of Egr1.
Error bars represent SEM. ns=not significant, *p<0.05, **p<0.01, ***p<0.001.

**Supplementary Figure 7.** Influence of Stabilin-ligands on Egr1 expression
Left: Schematic of BMDM treatment with Stabilin-ligands. Right: Relative Egr1 expression (RT-qPCR) of Stabilin-ligand treated BMDM. n=5.
Error bars represent SEM. ns=not significant, *p<0.05, **p<0.01, ***p<0.001.

**Supplementary Figure 8.** Plasma-treated BMDM and MAOEC
(**A**) Venn-Diagram showing differentially expressed genes (adjusted p<0.05) in ApoE-Stab1-KO (AS1, light red), ApoE-Stab2-KO (AS2, green), WT (blue) vs. ApoE plasma treated BMDMs. 4045 genes were significantly regulated comparing BMDM stimulated with ApoE-Stab1-KO plasma vs. ApoE-KO plasma (light red circle), 3918 genes were significantly regulated comparing BMDM stimulated with ApoE-Stab2-KO vs. ApoE-KO plasma (green circle), and 4121 genes were significantly regulated comparing BMDM stimulated with WT vs. ApoE-KO plasma (blue circle). 1714 genes were commonly dysregulated in BMDM stimulated with ApoE-Stab1-KO or ApoE-Stab2-KO vs. ApoE plasma (yellow circle). The majority of these commonly regulated

genes showed parallel upregulation or downregulation in both BMDM stimulated with ApoE-Stab1-KO plasma or ApoE-Stab2-KO plasma in comparison to ApoE-KO plasma (~70%). Genes that were significantly regulated in all three BMDM populations stimulated with WT, ApoE-Stab1-KO and ApoE-Stab2-KO plasma compared to ApoE-KO plasma (dark red circle) showed parallel upregulation or downregulation in ~76% of the 1230 genes (dark red circle).

**(B)** KEGG Pathways with common changes in ApoE-Stab1-KO, ApoE-Stab2-KO and WT in comparison to ApoE-KO. KEGG pathway analysis revealed that except for one all the pathways commonly altered in BMDM stimulated with plasma of WT, ApoE-Stab1-KO and ApoE-Stab2-KO mice as compared to BMDM stimulated with plasma of ApoE-KO mice showed gene alterations in the same direction as shown by NES.

**(C)** Relative Egr1 expression (RT-qPCR) in MAOEC stimulated with WT plasma, ApoE-KO plasma, ApoE-Stab1-KO plasma, ApoE-Stab2-KO plasma. n≥5.

Error bars represent SEM. ns=not significant, *p<0.05, **p<0.01, ***p<0.001.
